# An effector from the potato late blight pathogen bridges ENTH-domain protein TOL9a to an activated helper NLR to suppress immunity

**DOI:** 10.1101/2025.07.06.663370

**Authors:** Jogi Madhuprakash, AmirAli Toghani, Hsuan Pai, Madia Harvey, Adam R. Bentham, Benjamin A. Seager, Enoch Lok Him Yuen, Juan Carlos De la Concepción, David M. Lawson, Clare E. M. Stevenson, Angel Vergara-Cruces, Lida Derevnina, Tolga O. Bozkurt, Mark J. Banfield, Sophien Kamoun, Mauricio P. Contreras

**Affiliations:** The Sainsbury Laboratory, University of East Anglia, Norwich Research Park, Norwich, UK; Department of Plant Sciences, School of Life Sciences, University of Hyderabad, Gachibowli, Hyderabad, India; Department of Biochemistry and Metabolism, John Innes Centre, Norwich Research Park, Norwich, UK; Department of Life Sciences, Imperial College, London, UK

## Abstract

Pathogens counteract central nodes of NLR immune receptor networks to suppress immunity. However, the mechanisms by which pathogens hijack helper NLR pathways are poorly understood. Here, we show that an effector from the potato late blight pathogen *Phytophthora infestans* bridges the host protein NbTOL9a, a putative member of the host ESCRT pathway, to a helper NLR to suppress immunity. In this work, we solved the crystal structure of the RXLR-LWY effector AVRcap1b in complex with the ENTH domain of NbTOL9a. The structure revealed that unlike other RXLR-LWY effectors, AVRcap1b has a novel L-shaped fold that defines a new structural family of effectors in the *Phytophthora* genus. Moreover, we defined the AVRcap1b/NbTOL9a binding interface and designed effector mutants that don’t bind NbTOL9a, impairing immune suppression. This indicates that ENTH binding is required for full virulence activity of this effector. Lastly, we show that AVRcap1b associates specifically with activated NbNRC2 independently of NbTOL9a binding. This suggests that the effector functions as a bridge that interconnects NbNRC2 with the NbTOL9a pathway. These results illustrate an unprecedented pathogen mechanism to hijack helper NLR pathways and suppress immunity.

## Introduction

Immune receptors of the nucleotide binding and leucine-rich repeat (NLR) class are important components of innate immunity across all kingdoms of life (*1–4*). Upon recognition of pathogen-derived ligands, they subsequently initiate an array of immune responses to counteract infection. In plants, NLRs can be activated by pathogen-secreted virulence proteins, termed effectors, which pathogens deliver into host cells to modulate host physiology (*1*). A hallmark of NLR across eukaryotes and prokaryotes is their oligomerization into higher-order immune complexes, termed resistosomes in plants, or inflammasomes in bacteria and metazoans. These oligomeric complexes initiate immune signaling via diverse mechanisms, often leading to a form of programmed cell death (hypersensitive response) that hinders disease progression (*3, 5*). While effectors are typically thought of as activators of NLR-mediated immunity, an emerging concept is that many effectors can act as suppressors of NLR signaling (*1, 6–9*). Given the robust immunity mediated by NLRs following pathogen recognition, there is an intense evolutionary pressure to evolve effectors that can interfere with NLR activation. Indeed, several examples of NLR-suppressing effectors have been described to date (*1, 8*). However, despite growing insights into NLR activation, how pathogen effectors compromise NLR activation and NLR-mediated immunity to promote disease remains largely unknown.

NLRs belong to the signal adenosine triphosphatases (ATPases) with numerous domains (STAND) superfamily (*10*). They typically exhibit a tripartite domain architecture consisting of an N-terminal signaling domain, a central nucleotide binding domain, and C-terminal superstructure forming repeats (*1, 5, 10*). The central domain, termed NB-ARC (nucleotide binding adaptor shared by APAF-1, plant R proteins, and CED-4) in plant NLRs, is a hallmark of this protein family and plays a key role as a molecular switch, mediating conformational changes required for activation. In contrast, the N-terminal domain of plant NLRs is variable; it can broadly be used to classify these receptors into distinct groups, which tend to cluster together in NB-ARC–based phylogenetic analyses. In plants, coiled-coil (CC)–type and Toll–interleukin-1 receptor (TIR)–type N-terminal domains are the most widespread N-terminal domains. The N-terminal domains dictate the downstream immune signaling mechanisms of the resistosome oligomers, with CC-NLRs exhibiting calcium channel activity at the plasma membrane (*5, 10*).

There are diverse pathways of NLR activation and signaling. In some cases, one NLR protein, termed a functional singleton, can mediate both effector perception and subsequent immune signaling (*11*). However, some NLRs function together as receptor pairs or, in higher-order configurations, termed immune receptor networks (*1, 12, 13*). In these cases, one NLR acts as a pathogen sensor, requiring a second helper NLR to execute immune signaling. In Asterids, the largest group of flowering plants, the NRC immune receptor network is composed of multiple sensor NLRs and cell-surface receptors that require an array of downstream helper NLRs termed NRCs (NLRs required for cell death) to successfully initiate immune signaling (*14–16*). The NRC network can account for to about 90% of the NLRome in some plant species and plays a key role in immunity against a broad range of pathogens and pests including oomycetes, bacteria, viruses, nematodes, and insects, underscoring its agronomic importance (*6, 14, 17, 18*). Previously, we proposed that sensor-helper pairs in the NRC network function according to an activation-and-release model (*18*). Upon perception of their cognate effectors, activated sensors undergo conformational changes to expose their central NB-ARC domain and activate downstream NRC helpers via a transient interaction. This ultimately leads to NRC activation and immune signaling via the assembly of oligomeric resistosomes (*17–24*). We recently reported the cryo-EM structures of the resting state and activated helper NLR NbNRC2 from *Nicotiana benthamiana* (NbNRC2). These structures revealed that NbNRC2 accumulates as a homodimer before undergoing major conformational changes to convert into a homohexameric resistosome following activation by the virus resistance and NRC-dependent sensor NLR protein Rx (*21, 22*).

Given the central role of NLRs in plant immunity, it is not surprising that some parasite effectors have evolved to suppress NLR-mediated immunity. While some effector act indirectly by targeting host proteins downstream of NLR signaling, others inhibit immunity more directly by interacting with NLRs themselves (*6, 7, 25, 26*). In NLR networks, helper NLRs are critical nodes which integrate signals from sensor NLRs and cell-surface receptors and, as such, they represent optimal targets for pathogen effectors (*1, 8, 27*). Indeed, an emerging paradigm is that some pathogens compromise immunity mediated by NLR networks via effectors that bind helper NLRs (*6, 7*). For example, the potato cyst nematode (*Globodera rostochiensis*) effector SS15 can inhibit cell death mediated by *Nicotiana benthamiana* helper NLRs NbNRC2 and NbNRC3 by binding a hinge-like loop in the central NB-ARC domain of these NLRs, locking them in their autoinhibited resting state and preventing resistosome formation (*6, 9, 17*). We previously reported that the potato late blight pathogen, *Phytophthora infestans* deploys an effector AVRcap1b which can also suppress immune signaling and cell death mediated by NbNRC2 and NbNRC3. However, unlike SS15, AVRcap1b does not physically associate with resting state NbNRC2 or NbNRC3 and the biochemical mechanisms by which it suppresses these helper NLRs remain poorly understood (*6*).

*P. infestans* (oomycete, Peronosporales) is a major threat to potato (*Solanum tuberosum*) and tomato (*Solanum lycopersicum*) crops worldwide, and it also infects other Solanaceae, such as the model plant *N. benthamiana* (*28, 29*). Peronosporales species secrete a superfamily of effector proteins known as RXLR effectors; named after the conserved Arginine-any amino acid-Leucine-Arginine motif that follows the signal peptide of the proteins and is required for their translocation into host cells (*30, 31*). The *P. infestans* genome harbors about 550 predicted RXLR effectors that are grouped into ∼150 families and tend to exhibit sequence and expression polymorphisms between pathogen strains (*32–34*). Within the broader *Phytophthora* genus phylogeny, *P. infestans* belongs to clade 1c. Aside from *P. infestans* and *Phythophthora andina* —pathogens of the Solanaceae—this clade includes the host-specialized species *Phytophthora ipomoeae* and *Phytophthora mirabilis* that infect botanically distant plants in the Convolvulaceae and Caryophyllaceae families, respectively (*35, 36*). Previously, comparative analyses of effectors from different *Phytophthora* species have helped reveal how these effectors execute their virulence function and how these functions have evolved following host jumps (*36*).

The 678 amino acid effector AVRcap1b is a core *P. infestans* effector in the RXLR-LWY family (*37, 38*). Downstream of the signal peptide and RXLR motif, effectors in this family feature an effector domain composed of 1 WY domain and 1 or more LWY domains which typically consist of 5 α-helices (*30, 39, 40*). The best studied example in this family is the 670 amino acid Phytophthora suppressor of RNAi 2 (PSR2) effector from the soybean pathogen *Phytophthora sojae*. The crystal structure of PSR2 revealed a highly organized strand of WY/LWY repeats that are packed to form a rigid linear shape. Like PSR2, AVRcap1b features an effector domain composed of one WY domain and 6 tandem LWY domains. Whether AVRcap1b or other multi-repeat RXLR-LWY effectors also adopt a linear structure remains unclear to date.

In a previous study, we showed that AVRcap1b associates with the host plant protein NbTOL9a. AVRcap1b genetically requires NbTOL9a for full suppression of NRC-mediated cell death, suggesting that AVRcap1b is co-opting this host protein to suppress NLR-mediated signaling. NbTOL9a belongs to the Target of Myb 1-like (TOL) family of proteins, which are presumed to function as ubiquitinated cargo adaptors in the endosomal sorting complex required for transport (ESCRT) vesicle trafficking pathway (*41*). TOL proteins feature an N-terminal epsin N-terminal homology (ENTH) domain followed by a GGAs and Target of Myb 1 (GAT) domain, with these two domains being involved in binding to ubiquitinated plasma membrane (PM)-associated cargo. Following cargo binding, TOL proteins subsequently recruit the ESCRT-1 complex as well as other downstream components of the ESCRT pathway (*41, 42*). However, the mechanism by which AVRcap1b association with NbTOL9a leads to immune suppression is unknown.

In this study, we investigate the biochemical and structural mechanisms underlying AVRcap1b co-option of NbTOL9a and suppression of NRC-mediated signaling. We show that AVRcap1b directly binds the ENTH domain of NbTOL9a and we determine the crystal structure of the AVRcap1b-NbTOL9a ENTH complex, identifying a novel effector-host target interaction interface. The structure of AVRcap1b revealed a novel L-shaped fold distinct from previously characterized RXLR-LWY effectors. This L-shaped fold defines a new family of WY/LWY effectors that appear to be widespread across the *Phytophthora* genus. We demonstrate that AVRcap1b binding to NbTOL9a is required for the effector to fully suppress NbNRC2. Remarkably, although AVRcap1b does not interact with NbNRC2 in its resting state, it associates with sensor-activated NbNRC2 through an interface distinct from that used for NbTOL9a binding, likely forming a ternary complex. Overall, this work expands our understanding of the structural diversity of oomycete RXLR-LWY effectors and sheds light on the structural and biochemical mechanisms underpinning NLR suppression by pathogen effectors.

## Results

### AVRcap1b binds the ENTH domain of NbTOL9a

In a previous study, we identified NbTOL9a, a member of the TOL protein family, as a host target of AVRcap1b (*6*). We also showed that while AVRcap1b associates with NbTOL9a, it does not associate with other host TOL proteins such as NbTOL6. To gain further insights regarding AVRcap1b-TOL interactions, we leveraged the differential AVRcap1b association between NbTOL9a and NbTOL6 to identify the domain that determines TOL-AVRcap1b interactions. We generated a series of NbTOL9a-NbTOL6 chimeric proteins (**Fig. 1A**), which we subsequently assayed for AVRcap1b association via *in planta* co-immunoprecipitation. One chimeric variant of NbTOL6 carrying the N-terminal ENTH domain of NbTOL9a (NbTOL6^ENTH^) gained association with AVRcap1b (**Fig. 1B**).

**Figure 1:**
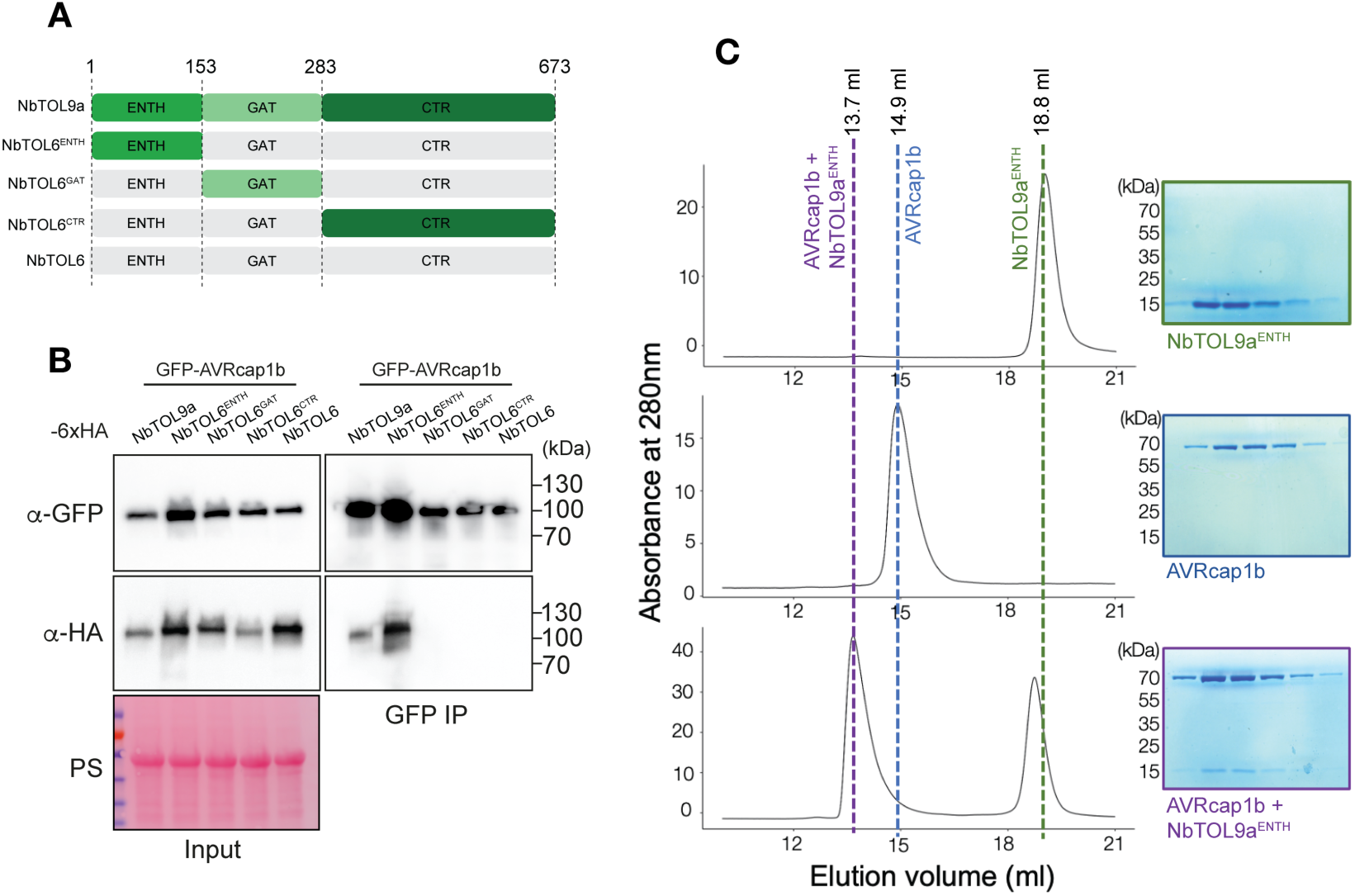
AVRcap1b binds the ENTH domain of NbTOL9a. (**A**) Schematic representation of chimeric TOL proteins used in co-immunoprecipitation (CoIP) experiments. Numbers indicate amino acid positions based on NbTOL9a numbering (**B**) CoIP experiments between AVRcap1b and three chimeric NbTOL proteins, NbTOL6^ENTH^, NbTOL6^GAT^ and NbTOL6^CTR^. NbTOL9a and NbTOL6 were included as positive and negative controls for AVRcap1b association, respectively. IPs were performed with agarose beads conjugated to GFP (GFP IP) antibodies. Total protein extracts were immunoblotted with appropriate antisera labelled on the left. Approximate molecular weights (kDa) of the proteins are shown on the right. RuBisCO loading controls were conducted using Ponceau staining. Experiments were independently repeated twice with similar results. (**C**) AVRcap1b binds the ENTH domain of NbTOL9a in vitro. Analytical size-exclusion chromatography (Superdex™ 200 10/300 GL column) was performed with AVRcap1b alone (top), NbTOL9a^ENTH^ alone (middle), or a 1:1 protein mixture (bottom). Elution profiles are shown with corresponding SDS-PAGE analyses of collected fractions. Approximate molecular weights (kDa) are indicated on the left of the SDS page gels.

To further characterize this interaction, we expressed and purified AVRcap1b and an NbTOL9a truncation consisting of only the ENTH domain (NbTOL9a^ENTH^), using *Escherichia coli* as a heterologous expression system. We successfully obtained homogeneous preparations of AVRcap1b and the NbTOL9a^ENTH^ but were unable to purify full-length NbTOL9a due to poor solubility and stability. Size exclusion chromatography analysis revealed that AVRcap1b and NbTOL9a^ENTH^ eluted at 14.9 and 18.8 mL, respectively (**Fig. 1C**). To assess complex formation *in vitro* we mixed the individually purified AVRcap1b and NbTOL9a^ENTH^ proteins and subjected the mixture to size exclusion chromatography. This resulted in a peak shift with an elution volume of 13.7 mL, consistent with complex formation. SDS-PAGE assessment of the peak fractions confirmed the presence of both proteins (**Fig. 1C**). To independently validate this interaction, we performed SEC-MALS with purified AVRcap1b and NbTOL9a^ENTH^, which orthogonally corroborated complex formation *in vitro* (**Fig. S1**). These results demonstrate that AVRcap1b directly interacts with NbTOL9a via its N-terminal ENTH domain, and that this domain alone is sufficient for the interaction.

### The crystal structure of AVRcap1b in complex with the NbTOL9a ENTH domain reveals a novel L-shaped effector fold

To structurally characterize AVRcap1b and define its interface with NbTOL9a, we crystallized the mature AVRcap1b protein (amino acids 62-678) in complex with NbTOL9a^ENTH^. We solved the structure using X-ray diffraction to a resolution of 4.1 Å (**Fig. 2A**, **Table S1**). AVRcap1b features an N-terminal secretion signal and an RXLR-DEER motif (amino acids 1-61) followed by a WY module (WY1) and 6 LWY modules (LWY2-7) (**Fig. 2A**). The structure includes the full ENTH domain of NbTOL9a and residues 78 to 675 of AVRcap1b, spanning most of the WY1 domain and the entirety of LWY2 through LWY7, with the exception of the final three residues of LWY7. The crystal structure revealed that AVRcap1b binds the ENTH domain of NbTOL9a via its WY1 and LWY2 regions and adopts a previously uncharacterized L-shaped fold (**Fig. 2A**). This architecture is notably distinct from the linear configuration of *P. sojae* PSR2, a structurally characterized RXLR-LWY effector (**Fig. 2B**). Comparisons with PSR2 indicate that the bend in AVRcap1b arises from the LWY5 domain, which introduces a 77° angle in the protein backbone, producing the L-shaped conformation (**Fig. S2**). We compared the structures of individual LWYs (LWY2–LWY7) from AVRcap1b and PSR2 in a pairwise manner. Alignments of each LWY to AVRcap1b LWY5 consistently produced the highest RMSDs, with an average of 6.078 Å compared to other AVRcap1b LWYs, and 6.834 Å when compared to each PSR2 LWY. These findings indicate that LWY5 shares low structural similarity with the other repeats. In contrast, all pairwise comparisons that excluded LWY5 showed lower RMSDs, averaging less than 4.827 Å (**Table S2**). Together, these results suggest that LWY5 is structurally distinct from the other LWY repeats in AVRcap1b and from the LWY domains of PSR2 and likely plays a key role in establishing the unique L-shaped arrangement observed in AVRcap1b (**Fig. S2. Table S2**).

**Figure 2:**
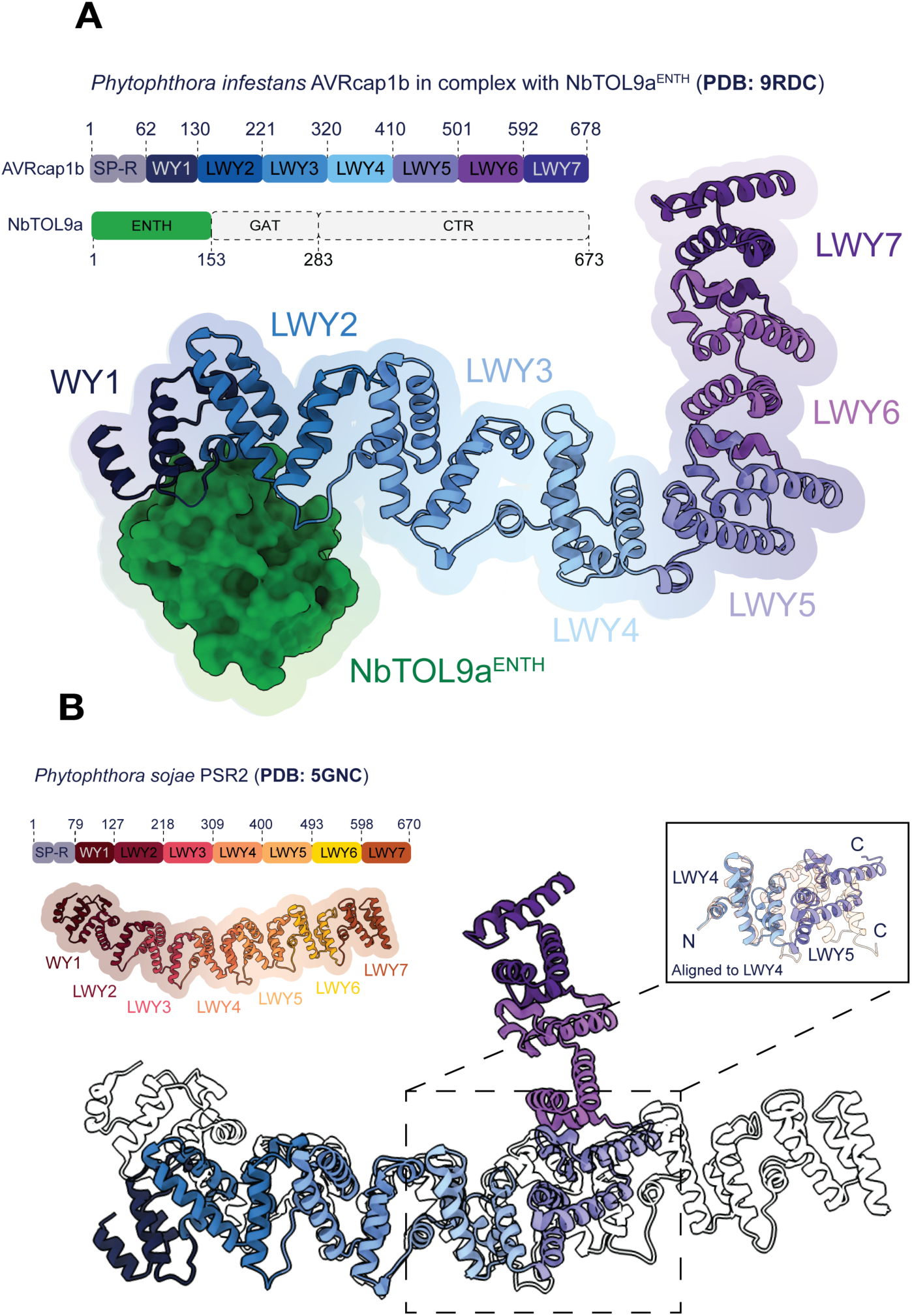
The crystal structure of AVRcap1b in complex with NbTOL9a^ENTH^ reveals an L-shaped fold. (**A**) Overall structure of *P. infestans* AVRcap1b bound to the ENTH domain of NbTOL9a (PDB ID: 9RDC). Individual structural repeat units of AVRcap1b and NbTOL9a ENTH domain are color-coded as indicated in the schematic above, which also denotes the amino acid boundaries for each repeat/domain. The schematic highlights the signal peptide (SP) and RXLR-DEER motif (R) at the N-terminus. See **Table S1** for a summary of data collection and processing statistics. (**B**) Structural superimposition of *P. infestans* AVRcap1b with the previously solved RXLR-LWY effector PSR2 from *P. sojae* (PDB ID: 5GNC). Domain boundaries of PSR2 are indicated in the schematic (top left). The inset (top right) shows a comparison of LWY repeats 4 and 5 from both effectors, aligned to LWY4, highlighting the divergence in overall conformation.

He et al. (*37*) previously proposed that the linear arrangement of LWY repeats in PSR2 is maintained by a conserved feature known as Loop^4-5^, connecting α4 and α5 helices in each LWY domain, which interacts with a hydrophobic pocket formed by four conserved leucine residues in the subsequent LWY unit. Given the L-shaped fold of AVRcap1b, we investigated whether this feature is conserved in this effector. We performed multiple sequence alignment of all AVRcap1b LWY domains, focusing on the conservation of Loop^4-5^ and the hydrophobic pocket it interacts with (**Fig. S3**). Interestingly, we observed that both Loop^4-5^ and the four conserved hydrophobic residues are conserved in LWY2 through LWY7 of AVRcap1b. This suggests that the L-shaped conformation of AVRcap1b is not due to the loss of these conserved structural features and that the presence of Loop^4-5^ and its cognate hydrophobic pocket is not sufficient for establishing a linear arrangement in RXLR-LWY effectors.

### AVRcap1b defines a novel family of L-shaped LWY effectors found across the Phytophthora genus

Given the novel L-shaped structure of AVRcap1b, we investigated the structural diversity of its orthologs and other members of the family. First, we performed a BLAST sequence similarity search against the NCBI non-redundant protein database using the AVRcap1b sequence as the query, retaining the top 250 hits (**Data S1**) (*43*). After filtering for sequences of similar length to AVRcap1b, we obtained a final dataset of 180 sequences from 18 Phytophthora species spanning clades 1, 2, 3, 4, 7, 8, and 11 (**Fig. 3A, Data S2**) (*44, 45*). Notably, PSR2 was among the hits. We then carried out phylogenetic analysis on the 180 sequences, supplemented with *P. infestans* AVRcap1b and three orthologs from *Phytophthora andina*, *Phytophthora mirabilis*, and *Phytophthora ipomoeae* (*35*) (**Data S3**). The resulting tree revealed two distinct, well-supported phylogenetic clades: one containing 30 sequences clustering with PSR2 and the other comprising 152 sequences clustering with AVRcap1b (**Fig. 3A**). To explore structural variation, we selected representative sequences from each clade across a range of *Phytophthora* species and taxonomic groups (**Fig. 3A**), and predicted their structures using AlphaFold 3 (*46*). We also modeled AVRcap1b and PSR2 as controls. Both AVRcap1b and PSR2 were predicted with high confidence (pTM scores of 0.72 and 0.67, respectively) and aligned closely with their corresponding crystal structures (RMSDs of 1.051 Å and 1.101 Å, respectively; **Fig. 3B**).

**Figure 3.**
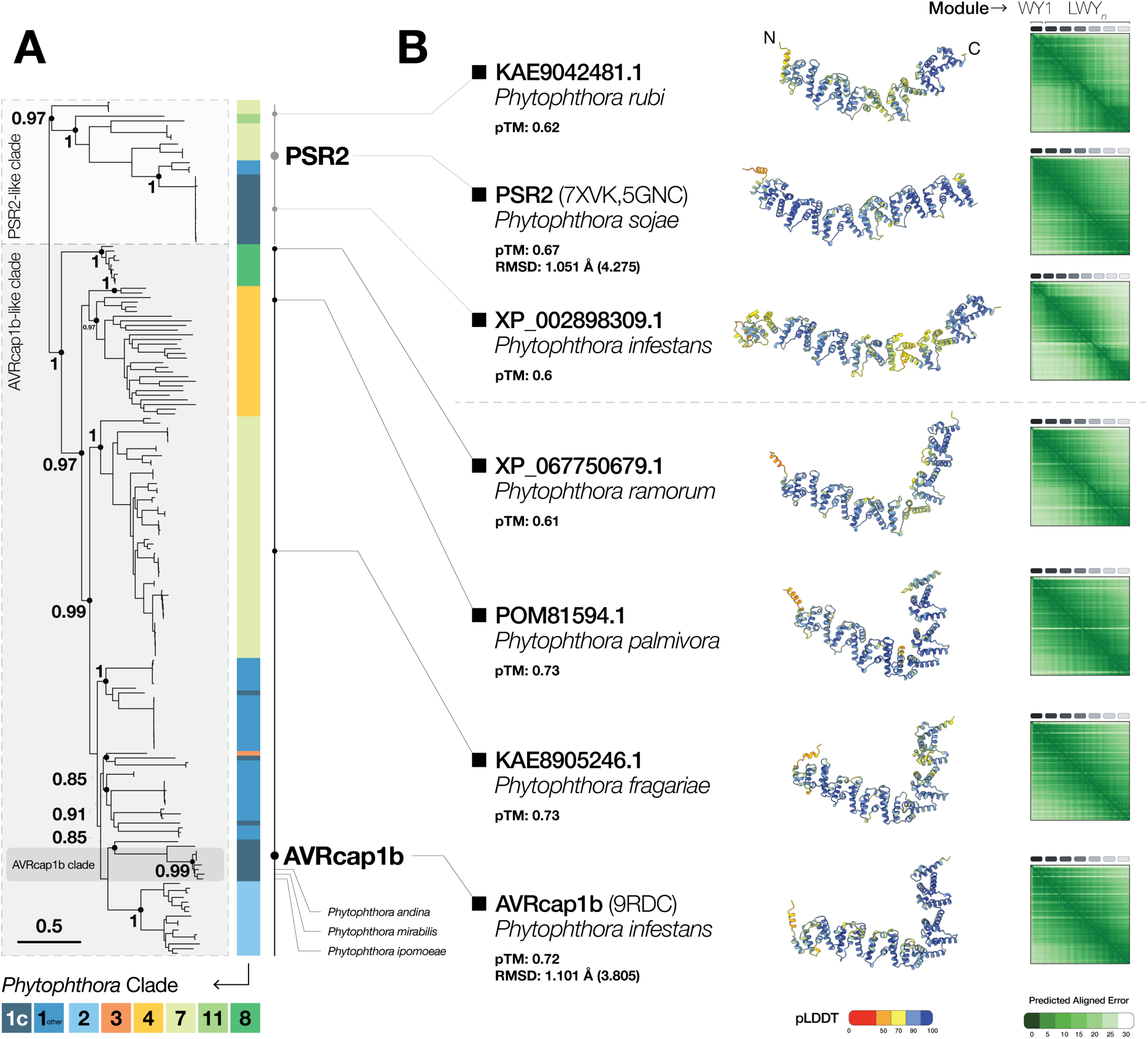
AVRcap1b defines a novel family of L-shaped LWY effectors found across the *Phytophthora* genus. (**A**) Phylogenetic tree of 184 LWY effectors identified by a PSI-BLAST search using AVRcap1b as the query against the NCBI non-redundant protein database. The tree resolves into two well-supported clades: one containing PSR2-like sequences and the other AVRcap1b-like sequences. Sequences are annotated by their Phytophthora clade of origin. The AVRcap1b subclade, which includes orthologs from other clade 1c species such as *P. andina*, *P. mirabilis*, and *P. ipomoeae*, is highlighted in dark gray. Node values represent bootstrap support from FastTree v2.1.11 (*47*). (**B**) AlphaFold 3-predicted structures of representative sequences from the PSR2-like and AVRcap1b-like clades. All structures were predicted with high confidence (pTM > 0.6), and models of PSR2 and AVRcap1b closely matched their respective crystal structures with RMSD values of 1.051 Å and 1.101 Å, respectively. Values in parentheses indicate unpruned RMSD values. PSR2-like sequences exhibit a stick-like topology, whereas AVRcap1b-like sequences consistently display an L-shaped fold across phylogenetically diverse clades.

Strikingly, all tested members of the AVRcap1b clade consistently exhibited an L-shaped topology, whereas all PSR2-like sequences exhibited stick-like structures (**Fig. 3B**). To further examine the AVRcap1b clade, we modeled its close orthologs—all from *Phytophthora* clade 1c—using AlphaFold 3. These orthologs consistently yielded high-confidence L-shaped models (pTM ≥ 0.65; **Fig. S4**) One exception was a truncated ortholog from *P. infestans* (XP_002896947.1), which lacked the LWY6 and LWY7 repeats and exhibited a shorter stick-like structure (**Fig. S4**). Overall, our results suggest that the L-shaped effector fold is conserved and widely distributed across phylogenetically distant *Phytophthora* clades.

### AVRcap1b binding interface in the NbTOL9a ENTH domain is polymorphic across NbTOL proteins

Interface analysis of the AVRcap1b-NbTOL9a^ENTH^ co-crystal structure revealed that the N-terminal WY1 and LWY2 domains of AVRcap1b engage the NbTOL9a ENTH domain (**Fig. 1A**, **Fig. 4A**) This interaction buries 791 Å^2^ of surface area on AVRcap1b and is stabilized by the insertion of α1 and α2 helices of NbTOL9a^ENTH^ into a shallow concave pocket formed by WY1 and LWY2. We identified several AVRcap1b residues within this interface involved in electrostatic and hydrogen-bond interactions that likely contribute to complex stability (**Fig. 4A** and **Fig. 4B**). Within WY1, R90 (located in the loop between α2 and α3), along with P92, G94, and K98 from α3, participate in binding. From LWY2, S139 and S143 (both from α1) also contribute to the interaction. These residues form hydrogen bonds with NbTOL9a^ENTH^ residues M5, R8, L14, and I15 from α1, and D18, A20, M21, D24, D27, and I28 from α2.

**Figure 4:**
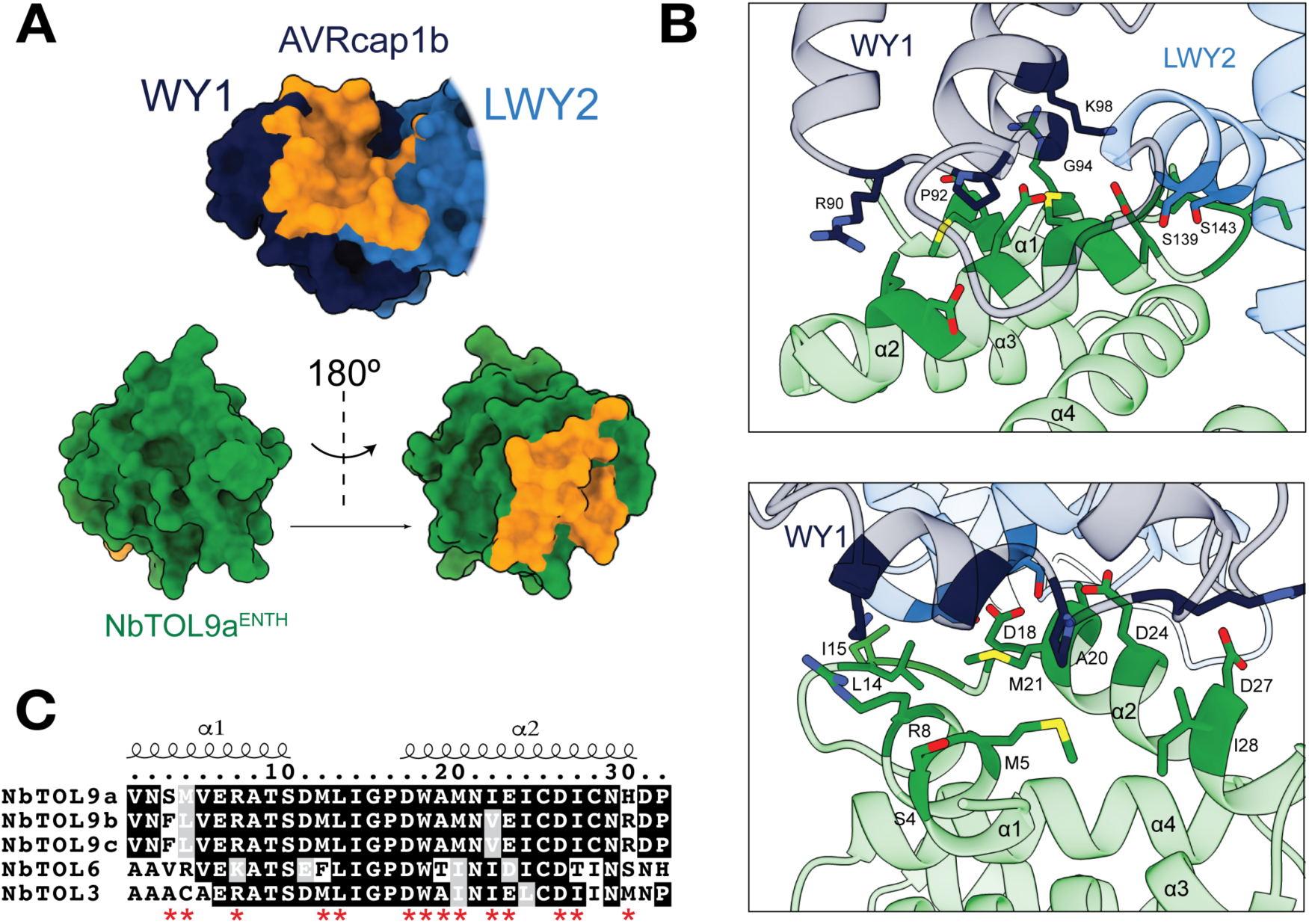
AVRcap1b-NbTOL9a^ENTH^ binding interface reveals key residues involved in the interaction. **(A)** Interaction interface between AVRcap1b and NbTOL9a ENTH domain truncation (NbTOL9a^ENTH^). Surfaces involved in the interaction are shown in orange. (**B**) Insets show the interaction interface between AVRcap1b and NbTOL9a^ENTH^. Potential contact residues from all these interfaces are highlighted in stick representation. (**C**) Amino acid sequence alignment of all 5 *N. benthamiana* TOL proteins, focusing on the AVRcap1b interaction interface. Positions of key residues involved in the interaction are highlighted with a red asterisk. Secondary structure and amino acid numbering (based on NbTOL9a) is shown at the top.

We previously reported that AVRcap1b strongly associates with NbTOL9a, exhibits weak association with NbTOL9b and NbTOL9c, and does not bind NbTOL6 or NbTOL3. Sequence comparison of the ENTH domains revealed polymorphisms at key interacting residues in these paralogs, which likely account for the reduced or absent binding to AVRcap1b (**Fig. 4C**).

### Mutagenesis of the interaction interface compromises NbTOL9a binding and AVRcap1b-mediated suppression of NbNRC2

To experimentally validate the AVRcap1b-NbTOL9a^ENTH^ co-crystal structure and to assess the functional relevance of this interaction interface in the suppression of NRC-mediated immunity, we generated six AVRcap1b point mutants targeting residues at the ENTH-binding interface within WY1 and LWY2 modules. Each of the six interface residues within WY1 and LWY2 modules (**Fig. 4B**) was substituted with glutamic acid, and the resulting variants were tested for NbTOL9a association via *in planta* co-immunoprecipitation. Four variants, AVRcap1b^P92E^, AVRcap1b^G94E^, AVRcap1b^S139E^, and AVRcap1b^S143E^, lost any detectable interaction with NbTOL9a, consistent with their contribution to the crystal structure interface (**Fig. 5A**). One mutant, AVRcap1b^R90E^, exhibited drastically reduced NbTOL9a association. These findings support the importance of these residues for NbTOL9a binding (**Fig. 5A**).

**Figure 5:**
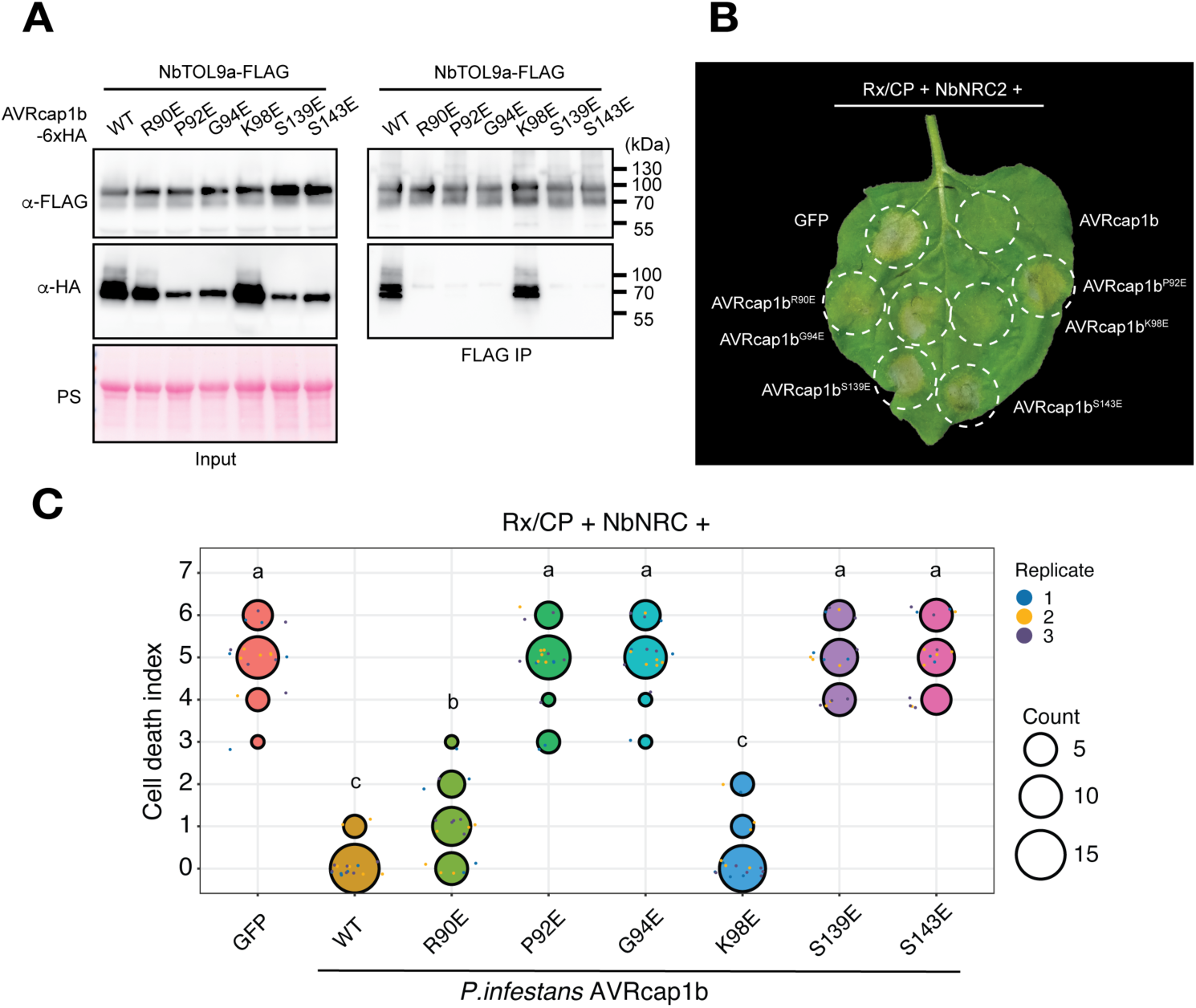
Mutations in the NbTOL9a-binding interface compromise AVRcap1b-mediated suppression of NbNRC2. (**A**) CoIP experiments between different AVRcap1b variants and NbTOL9a. IPs were performed with agarose beads conjugated to FLAG (FLAG IP) antibodies. Total protein extracts were immunoblotted with appropriate antisera labelled on the left. Approximate molecular weights (kDa) of the proteins are shown on the right. Rubisco loading controls were visualized by Ponceau staining. The experiment was independently repeated three times with similar results. (**B**) Photo of representative WT *N. benthamiana* leaf showing cell death after co-expression of Rx, PVX CP, NbNRC2 and different AVRcap1b variants in *nrc2/3/4* KO *N. benthamiana* plants. WT *P. infestans* AVRcap1b and GFP were included as positive and negative controls for suppression respectively. The experiments were repeated three times with at least 6 technical replicates per repeat, with similar results in all cases (**C**) Quantitative analysis of cell death assays shown in (**B**). Cell death was scored based on a modified 0-7 scale at 5-7 days post agroinfiltration (*48*). Results are presented as dot plots, where the size of each dot is proportional to the number of samples with the same score (Count). Data represent three biological replicates. Statistical analysis was carried out in R, using the BestHR package (*49*). Letters (“a”, “b” or “c”) above each treatment indicate statistically significant differences relative to the negative control (GFP).

We previously showed that AVRcap1b genetically requires NbTOL9a to fully suppress NRC2-mediated hypersensitive cell death (*6*). To investigate how disruption of this interaction compromises effector suppression activity, we assayed the six variants described above for their ability to suppress NbNRC2-mediated cell death triggered by Rx/CP activation in the *N. benthamiana* co-expression system. Notably, the same four variants that lost association to NbTOL9a were compromised in their capacity to suppress NbNRC2-mediated cell death (**Fig. 5B** and **Fig. 5C**). The R90E variant which exhibited weak association to NbTOL9a was still capable of suppressing NbNRC2-mediated cell, although not to the same extent as WT AVRcap1b (**Fig. 5B** and **Fig. 5C**). These results indicate that AVRcap1b binding to NbTOL9a is required for full suppression of NbNRC2-mediated immunity and confirm the functional relevance of the interaction surface identified in the co-crystal structure.

### NbTOL9a interaction interfaces show variability in different AVRcap1b clades

To investigate the conservation of the NbTOL9a ENTH domain-binding interfaces identified in *P. infestans* AVRcap1b among its orthologs, we analyzed the conservation of these sequences in the AVRcap1b-like proteins identified in the phylogenetic analysis (**Fig. 3**). Based on well-supported branches and taxonomic clustering, we grouped the AVRcap1b clade sequences into seven subclades (**Fig. 6**). We then generated sequence logo plots to examine conservation at the two key NbTOL9a ENTH domain interaction interfaces, located in the WY1 and LWY2 domains of AVRcap1b. Both interfaces exhibited substantial variability within and between clades, particularly at key contact residues. Notably, in clade 6—which includes AVRcap1b orthologs from *Phytophthora* clade 1c—of the five key interface residues, P92 was conserved across all non-truncated sequences (**Fig. 6**).

**Figure 6.**
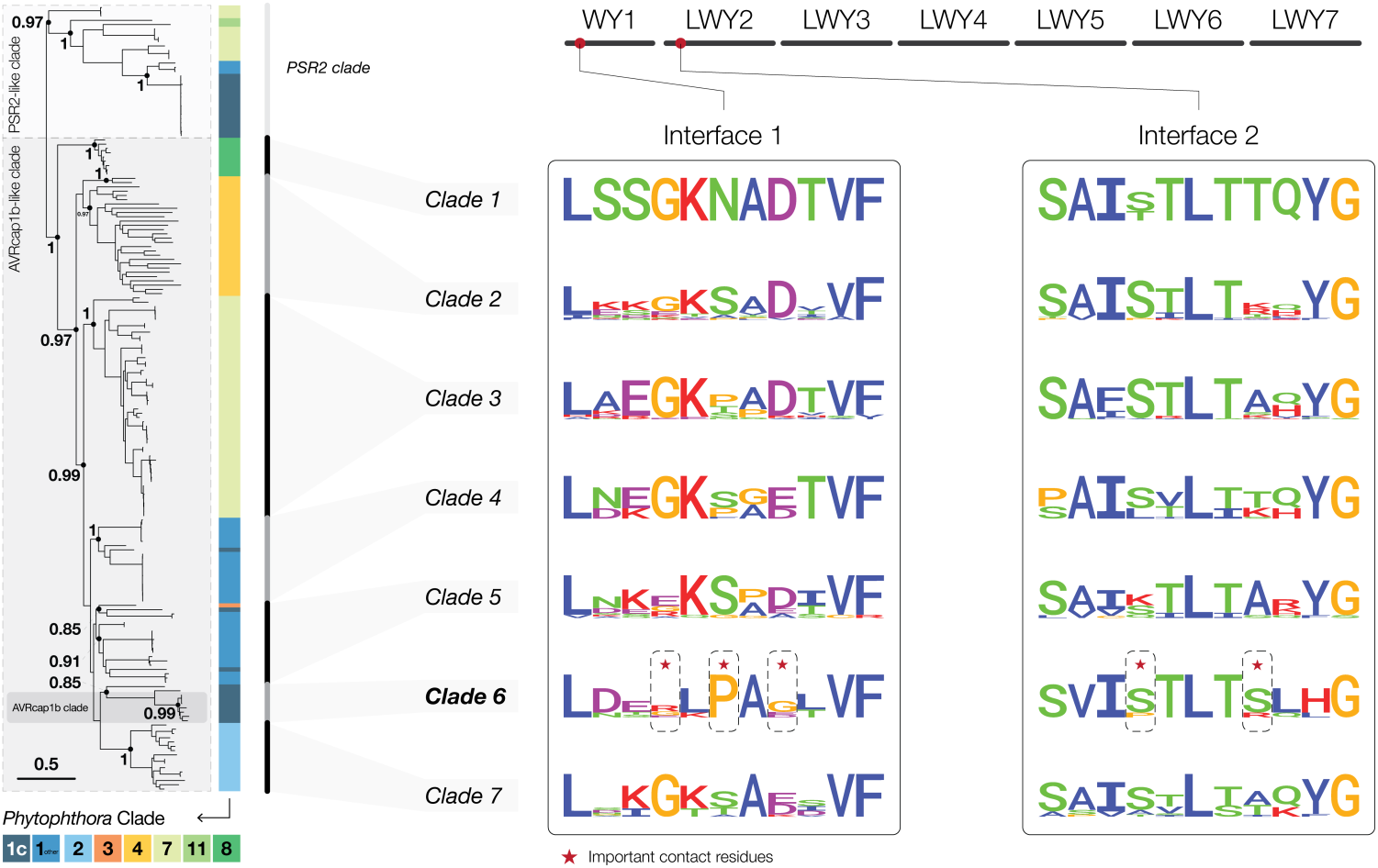
NbTOL9a interaction interfaces show variability across AVRcap1b clades. The AVRcap1b-like clade was divided into seven subclades based on well-supported phylogenetic branches and Phytophthora clade annotations. Sequence logo plots were generated for two NbTOL9a ENTH domain binding interfaces on AVRcap1b, located on the WY1 and LWY2 repeats, respectively, across all subclades. Both interfaces exhibited variability within and between subclades. In clade 6, which includes AVRcap1b and its clade 1c orthologs, only one of the five key contact residues—corresponding to P92 (interface 1)—was conserved across all sequences.

To support these observations, we performed Shannon entropy analysis on the full-length AVRcap1b-like sequences (**Data S4**). Shannon entropy quantifies amino acid variability at each alignment position, ranging from 0 (log₂1) when a position is fully conserved, to 4.32 (log₂20) when all 20 amino acids occur with equal frequency. Following previous studies, we defined positions with entropy >1.5 as highly variable (*16, 21, 50*). The number of highly variable sites fluctuates widely across clades, from 4 in clade 1 to 368 in clade 2, and these sites were distributed throughout the protein, often with clade-specific patterns (**Fig. S5**, **Table S3**). In clade 6, we identified 56 variable sites which surpassed the 1.5 threshold, out of which 10 were on WY1 and 5 on LWY2. While residues corresponding to R90 (entropy = 1.84) and G94 (entropy = 1.66) surpassed the threshold, other key contact points such as P92 (entropy = 0.59), S139 (entropy = 1.15), and S143 (entropy = 1.15) all had entropy values below 1.5 (**Table S3**). Together, these analyses reveal that the NbTOL9a-binding interfaces of AVRcap1b-like proteins are variable across clades, with some contact sites exhibiting both inter- and intra-clade diversity.

### Clade 1c AVRcap1b orthologs do not suppress NRC activity

To further investigate the link between structure and function in AVRcap1b, we leveraged the sequence diversity within *Phytophthora* clade 1c. Although we previously established that the NbTOL9a ENTH binding interface is overall variable across AVRcap1b clades, we examined its conservation across more closely related orthologs from *Phytophthora* clade 1c species identified in our phylogenetic analysis (**Figure 6**). All five key AVRcap1b residues involved in NbTOL9a ENTH domain binding, P92, G94, K98, S139, and S143, are conserved across these orthologs. R90 is retained in *P. infestans*, *P. ipomoeae,* and *P. andina*, whereas *P. mirabilis* carries a glycine at the equivalent position. These findings suggest that NbTOL9a binding is likely conserved among clade 1c AVRcap1b orthologs.

Next, we tested the degree to which AVRcap1b orthologs from clade 1c *Phytophthora* species (*P.mirabilis*, *P. ipomoeae*, and *P. andina*) suppress NbNRC2-mediated cell death following activation with Rx/CP. In addition to preserving all six key NbTOL9a ENTH domain-binding residues, these orthologs share high overall sequence similarity with the *P. infestans* AVRcap1b, with amino acid sequence identities of 88.5%, 90%, and 91%, respectively. Of the tested proteins, only *P. infestans* AVRcap1b suppressed NbNRC2-mediated responses, despite comparable expression levels for all tested proteins in *N. benthamiana* (**Fig. 7A, 7B**).

**Figure 7:**
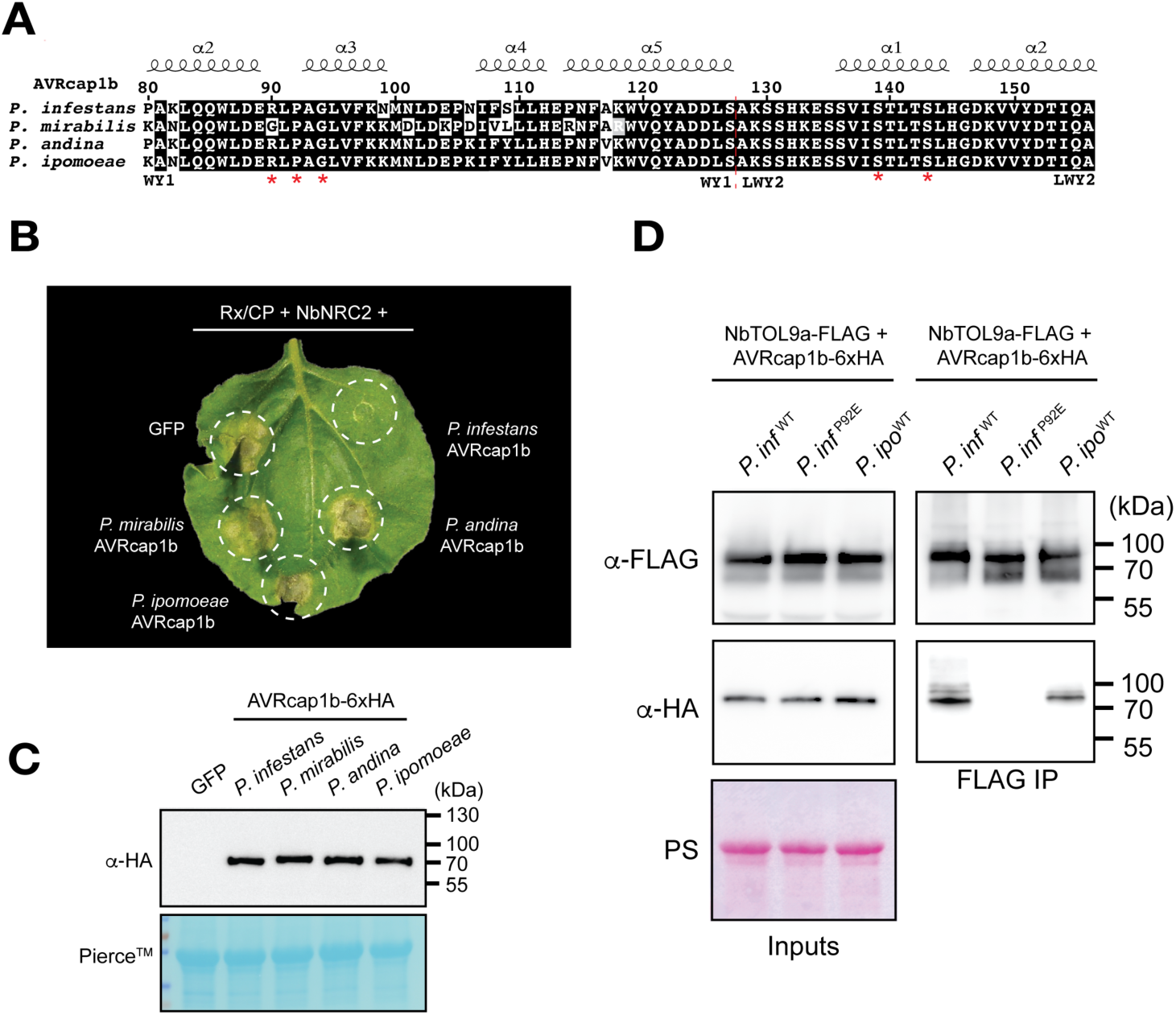
Despite conservation of NbTOL9a binding interface, AVRcap1b orthologs from *Phytophthora* clade 1c species do not to suppress NbNRC2. (**A**) Amino acid sequence alignment of AVRcap1b homologs from the four clade 1c *Phytophthora* species. Red asterisks denote critical residues for NbTOL9a binding. (**B**) Photo of representative WT *N. benthamiana* leaf showing cell death after co-expression of Rx/CP and NbNRC2 and Phytophthora clade 1c AVRcap1b orthologs in *nrc2/3/4* KO *N. benthamiana* plants. *P. infestans* AVRcap1b and GFP were included as positive and negative controls for suppression respectively. Cell death was photographed 5 days after agroinfiltration. (**C**) Expression of Phytophthora clade 1c AVRcap1b homologs was tested by Western blotting. GFP was included as a control for non-specific signal. Blots were probed with appropriate antisera (labeled on the left). Loading control was probed with Pierce Stain. Approximate molecular weights are displayed in kDa on the right. (**D**) Co-immunoprecipitation assays between NbTOL9a and AVRcap1b variants. C-terminally 3xFLAG-tagged NbTOL9a was co-expressed with C-terminally 6xHA-tagged AVRcap1b variants. IPs were performed with agarose beads conjugated to FLAG antibodies (FLAG IP). Total protein extracts were immunoblotted with the antisera labeled on the left. Approximate molecular weights (in kilodalton) of the proteins are shown on the right. Rubisco loading control was carried out using Ponceau stain (PS). The experiment was repeated three times with similar results.

We investigated NbTOL9a binding using *in planta* co-immunoprecipitation with the *P. ipomoeae* AVRcap1b ortholog. This ortholog associated with NbTOL9a at levels similar to *P. infestans* AVRcap1b, unlike the negative control AVRcap1b^P92E^ (**Fig. 7C**). These results indicate that while NbTOL9a binding is required, it is not sufficient for suppression of NbNRC2-mediated cell death. Additional determinants, potentially besides the conserved NbTOL9a binding interface, likely account for the effector immunosuppression activity.

### Clade 1c AVRcap1b orthologs do not suppress NRC activity

To further investigate the link between structure and function in AVRcap1b, we leveraged the sequence diversity within *Phytophthora* clade 1c. Although we previously established that the NbTOL9a ENTH binding interface is overall variable across AVRcap1b clades, we examined its conservation across more closely related orthologs from *Phytophthora* clade 1c species identified in our phylogenetic analysis (**Figure 6**). All five key AVRcap1b residues involved in NbTOL9a ENTH domain binding, P92, G94, K98, S139, and S143, are conserved across these orthologs. R90 is retained in *P. infestans*, *P. ipomoeae,* and *P. andina*, whereas *P. mirabilis* carries a glycine at the equivalent position. These findings suggest that NbTOL9a binding is likely conserved among clade 1c AVRcap1b orthologs.

Next, we tested the degree to which AVRcap1b orthologs from clade 1c *Phytophthora* species (*P.mirabilis*, *P. ipomoeae*, and *P. andina*) suppress NbNRC2-mediated cell death following activation with Rx/CP. In addition to preserving all six key NbTOL9a ENTH domain-binding residues, these orthologs share high overall sequence similarity with the *P. infestans* AVRcap1b, with amino acid sequence identities of 88.5%, 90%, and 91%, respectively. Of the tested proteins, only *P. infestans* AVRcap1b suppressed NbNRC2-mediated responses, despite comparable expression levels for all tested proteins in *N. benthamiana* (**Fig. 7A, 7B**).

We investigated NbTOL9a binding using *in planta* co-immunoprecipitation with the *P. ipomoeae* AVRcap1b ortholog. This ortholog associated with NbTOL9a at levels similar to *P. infestans* AVRcap1b, unlike the negative control AVRcap1b^P92E^ (**Fig. 7C**). These results indicate that while NbTOL9a binding is required, it is not sufficient for suppression of NbNRC2-mediated cell death. Additional determinants, potentially besides the conserved NbTOL9a binding interface, likely account for the effector immunosuppression activity.

### AVRcap1b associates specifically with activated NbNRC2 independently of NbTOL9a binding

Our finding that NbTOL9a binding is not sufficient for AVRcap1b-mediated suppression of NbNRC2 prompted us to revisit its interaction with NRC proteins. Despite its ability to suppress NbNRC2- and NbNRC3-mediated cell death, AVRcap1b did not associate with either helper NLR in our previous assays. (*6*). However, at the time, we were limited to studying AVRcap1b interactions with resting-state NRC proteins, as investigating interactions with activated forms was technically challenging due to the rapid onset of cell death upon immune receptor activation. To address this issue, we leveraged NbNRC2^EEE^, an NRC variant carrying mutations in its N-terminal MADA motif that abolish cell death while preserving receptor activation and oligomerization (*18, 51*). We co-expressed AVRcap1b with NbNRC2^EEE^ in *N. benthamiana,* activating the system using Rx/CP or using Rx/GFP as a non-activated control. *In planta* co-immunoprecipitation experiments revealed that AVRcap1b specifically associates with activated NbNRC2^EEE^, but not with its resting state, suggesting that AVRcap1b associates with NbNRC2 only following activation by a matching NLR sensor (**Fig. 8**). This observation challenges our earlier interpretations of indirect suppression (*6*) and supports a model in which AVRcap1b directly associates with sensor-activated NRC proteins.

**Figure 8:**
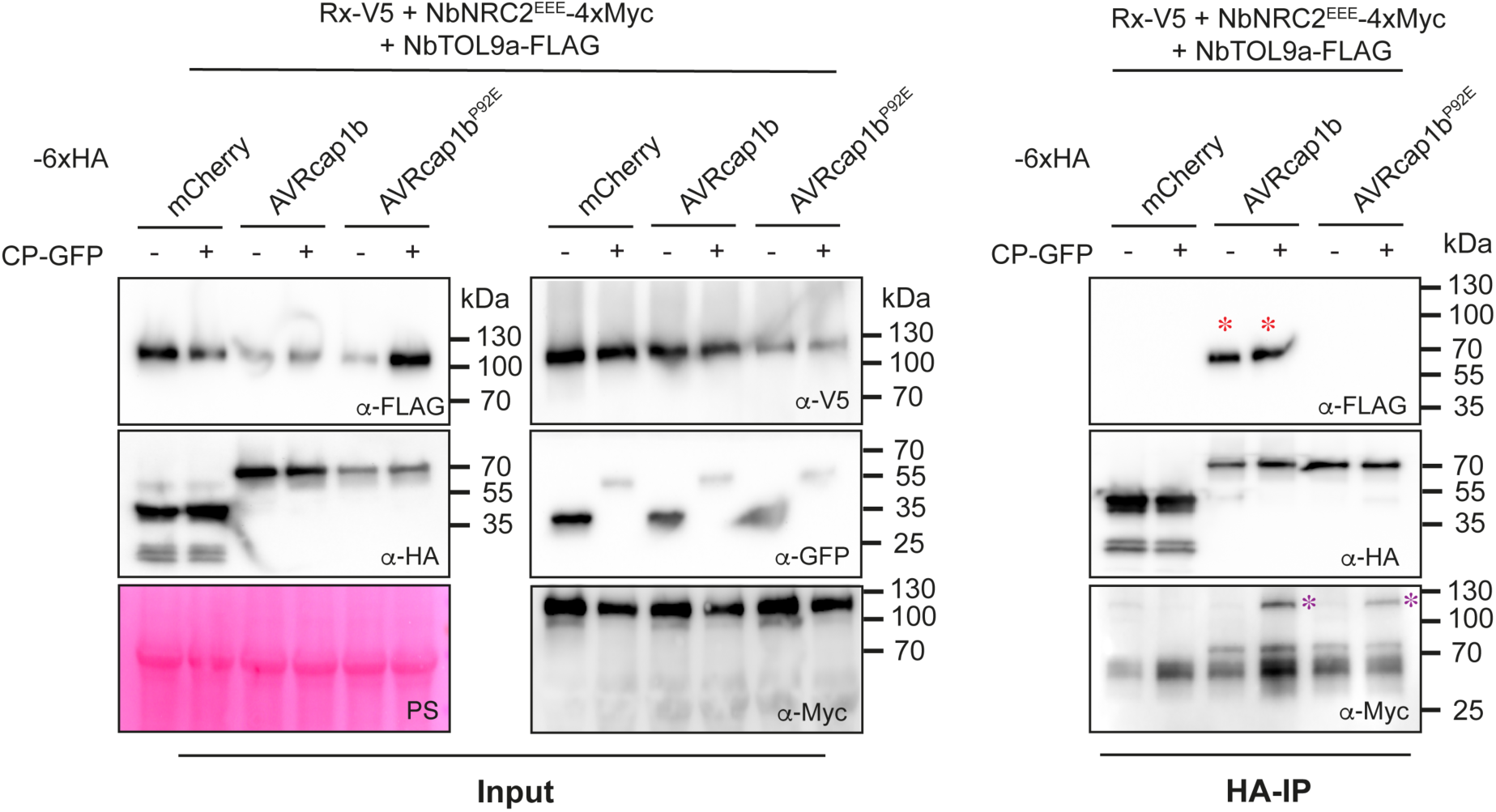
*P. infestans*AVRcap1b specifically associates with Rx/CP-activated NbNRC2. CoIP assays between resting state or activated NbNRC2 and AVRcap1b variants. C-terminally 3xFLAG-tagged NbNRC2 was co-expressed with C-terminally V5-tagged Rx and C-terminally 6xHA-tagged AVRcap1b. The Rx/NbNRC2 system was activated with C-terminally GFP tagged PVX CP. Free GFP used as a negative control for activation. IPs were performed with agarose beads conjugated to HA antibodies (HA IP). Total protein extracts were immunoblotted with the antisera labeled on the left. Approximate molecular weights (in kilodalton) of the proteins are shown on the right. Red asterisks indicate AVRcap1b association with NbTOL9a in both resting state and activated-NbNRC2 conditions. Purple asterisks indicate AVRcap1b and AVRcap1b^P92E^ association with Rx/CP activated NbNRC2^EEE^. RuBisCO loading control was carried out using Ponceau stain (PS). The experiment was repeated three times with similar results.

In these *in planta* co-immunoprecipitation experiments, we also probed for NbTOL9a following AVRcap1b pulldown. AVRcap1b associated at comparable levels with NbTOL9a under both non-activated (Rx/GFP) and activated (Rx/CP) conditions (**Fig. 8**), indicating that its association with NbTOL9a and activated NbNRC2 are not mutually exclusive and that the proteins likely form a ternary complex (**Fig. 8**). To further investigate this model, we performed co-immunoprecipitation assays with AVRcap1b^P92E^, a variant that disrupts NbTOL9a binding and compromises NbNRC2 suppression. As previously observed, AVRcap1b^P92E^ failed to associate with NbTOL9a but retained its specific association with activated NbNRC2^EEE^, mirroring the behaviour of the wild-type protein (**Fig. 8**).

Lastly, we tested the *P. ipomoeae* AVRcap1b ortholog in equivalent co-immunoprecipitation experiments with NbNRC2^EEE^ under resting state and sensor-activated conditions. Unlike *P. infestans* AVRcap1b, the *P. ipomoeae* ortholog did not associate with NbNRC2 under resting-state or activated conditions, even though it associated with NbTOL9a (**Fig. S6**). Overall, these results indicate that AVRcap1b associates with sensor-activated NbNRC2 via an interface distinct from that used to bind NbTOL9a, and that both NbTOL9a binding and interaction with activated NbNRC2 are required for AVRcap1b-mediated immunosuppression.

## Discussion

The aim of this study was to gain a better understanding of the molecular mechanism by which AVRcap1b binding to NbTOL9a results in suppression of NRC-mediated hypersensitive cell death. We found that AVRcap1b suppresses immunity by bridging the activated form of the helper NLR NbNRC2 and the host ESCRT pathway protein NbTOL9a. This model points to an unprecedented pathogen mechanism to hijack helper NLR pathways for immunosuppression. Our experiments revealed that AVRcap1b binds the ENTH domain of NbTOL9a to suppress NbNRC2-mediated signaling (**Fig. 1**, **Fig. 2**, **Fig. 4**, **Fig. 5**). The crystal structure of AVRcap1b-NbTOL9a^ENTH^ revealed that this effector exhibits a novel L-shaped fold, distinct from other RXLR-LWY effectors. Using phylogenomics and protein structure modelling, we show that L-shaped effectors are widespread throughout AVRcap1b clade proteins in the *Phytophthora* genus although they likely carry diverse activities given the lack of conservation of the NbTOL9a ENTH domain binding interface (**Fig. 3**). NbNRC2 suppression activity was specific to *P. infestans* AVRcap1b, with even the AVRcap1b ortholog from *P. ipomoeae* unable to suppress NbNRC2-mediated cell death, despite binding NbTOL9a (**Fig. 6**, **Fig. 7**). AVRcap1b, but not its *P. ipomoeae* ortholog, can associate with both NbTOL9a and sensor-activated NbNRC2 using distinct interfaces, consistent with a model in which AVRcap1b connects an activated helper NLR to NbTOL9a to suppress immunity (**Fig. 8**).

AVRcap1b defines a novel family of *Phytophthora* effectors with an L-shaped configuration, in contrast to PSR2 from *P. sojae*—the only other experimentally determined RXLR-LWY effector structure—which adopts a linear, stick-like shape. The linear PSR2 structure was previously proposed to be mediated by interactions between a conserved Loop^4-5^ and a hydrophobic patch in adjacent LWY modules (*37*). Interestingly, AVRcap1b features a bend introduced by LWY module 5, which disrupts this linearity despite conservation of Loop^4-5^ and the hydrophobic pockets in the LWY modules, suggesting that these features are not sufficient for a linear arrangement. The L-shaped fold appears to be widespread across effectors in the *Phytophthora* genus. Using AlphaFold 3, we predicted additional L-shaped LWY effectors across distantly related *Phytophthora* species that infect diverse plant hosts, suggesting that this is a conserved and widespread structural fold. Nonetheless, the precise contribution of this L-shaped arrangement to the immune suppression and virulence activities of AVRcap1b and other structurally related effectors remains unclear. Our computational analyses predict that most L-fold RXLR-LWY effectors do not bind NbTOL9a. Nonetheless, we cannot rule out the possibility that AVRcap1b-clade effectors interact with ENTH domain proteins from their respective host plants.

Structure-guided mutagenesis of the AVRcap1b-NbTOL9a^ENTH^ interface revealed that direct interaction between these two proteins is required for full NbNRC2 immunosuppression activity. TOL proteins, conserved across all eukaryotes, function as ubiquitin receptors in the ESCRT pathway (*41, 42*). Our previous work indicated that NbTOL9a acts as a negative regulator of NbNRC2 and NbNRC3-dependent cell death independently of AVRcap1b (*6*). Interestingly, ESCRT components have been shown to negatively regulate programmed cell death, by removing pore-forming proteins from membranes (*52–55*). Although the precise role of plant TOLs in membrane trafficking and remodelling remains unclear, our data together with previous findings support a model in which AVRcap1b exploits this negative regulatory mechanism to suppress immune activation. Whether AVRcap1b-mediated suppression of NRCs requires canonical ESCRT machinery or instead relies on ESCRT-independent functions of NbTOL9a remains to be determined. Importantly, clarifying the interplay between AVRcap1b, TOLs, and NLR-mediated cell death may reveal novel immune regulatory mechanisms and shed light on how these are exploited by effectors.

Interestingly, although TOL-binding is required for full immunosuppression by AVRcap1b, it is not sufficient. None of the clade 1c orthologs of *P. infestans* AVRcap1b were capable of suppressing NbNRC2, even though the NbTOL9a-binding interface is highly conserved across these proteins, and the *P. ipomoeae* ortholog can associate with NbTOL9a *in planta*. It remains possible that these AVRcap1b orthologs have also co-opted TOL proteins from their respective hosts, and it would be interesting to determine the degree to which they bridge such TOL proteins to NLRs or other host components to promote disease. In the future, functional and structural dissection of L-shaped LWY effectors from other *Phytophthora* species will advance our understanding of host target diversity and virulence mechanisms. Moreover, it will be interesting to determine the extent to which modulation of host TOL proteins by effectors is a conserved virulence strategy across the *Phytophthora* genus and other pathogens and pests.

Activated NRCs transition from resting state homodimers to hexameric resistosomes (*18*). Although we previously reported that AVRcap1b does not associate with the homodimeric resting state NbNRC2 (*6*), in this study we show that it can specifically associate with sensor-activated NbNRC2. However, it remains to be determined whether AVRcap1b blocks resistosome assembly at intermediate stages of activation or targets the mature hexameric resistosome. Although the structures of both resting state and activated NbNRC2 have been described (*21, 22*), the precise dynamics of NLR resistosome assembly and the structure of any assembly intermediates remain unclear. Further work will clarify the precise mechanisms by which AVRcap1b-NRC association leads to immune suppression.

We previously reported that nematode and oomycete effectors proteins have convergently evolved to counteract the NRC central nodes of a major NLR immune receptor network of the Solanaceae through different mechanisms (*6*). The potato cyst nematode effector SPRYSEC 15 (SS15) directly inhibits signaling of NRC1, NRC2 and NRC3 paralogs from diverse solanaceous species (*6, 9*). Structural analysis of SS15-NRC complexes revealed that SS15 binds NRCs and locks them in their autoinhibited resting state, likely preventing the conformational changes required for activation (*17*). Temporal dynamics of NRC activation and suppression must be critical to the outcome of the interaction. SS15 may be unable to suppress immune signaling if NRC activation has already occurred prior to its delivery into the host cell. In contrast, AVRcap1b may be capable of effectively suppressing NRCs even at later time points following activation. This could represent a more effective suppression mechanism, as AVRcap1b can inhibit NRC-mediated immune signaling even after recognition of the pathogen AVR effectors and activation of the sensor/helper NLRs. Further work will shed light on the precise interplay between NRC activation dynamics and AVRcap1b effector delivery and immunosuppression dynamics.

## Materials and Methods

### Plant growth conditions

Wild-type and *nrc2/3/4* CRISPR mutant (*56*) *N*. *benthamiana* lines were grown in a controlled environment growth chamber with a temperature range of 22 to 25°C, humidity of 45% to 65% and a 16/8-h light/dark cycle.

### Plasmid construction

We used the Golden Gate Modular Cloning (MoClo) kit (*57, 58*) and the MoClo plants part kit (*57, 58*) for cloning all constructs used for in planta expression. All vectors used were generated with these kits unless otherwise stated. Cloning design and sequence analysis were done using Geneious Prime (v2021.2.2; https://www.geneious.com). All AVRcap1b constructs used for in planta expression were cloned into the pICH86988 level one acceptor with a C-terminal 3xFLAG tag (pICSL5007) or a C-terminal 6xHA tag (pICSL5009). NbTOL9a, NbNRC2, Rx, PVX CP and NbTOL9a constructs were cloned into pJK001c-mRFP acceptor with 2x35s promoter (pICSL51288) and terminator (pICSL41414) as well as the C-terminal tag indicated. C-terminal tags used were 3xFLAG (pICSL5007), 6xHA (pICSL5009), 4xMyc (pICSL5010), V5 (pICSL5012) and eGFP (pICSL5034).

### Cell death assays by agroinfiltration in *Nicotiana benthamiana*

Proteins of interest were transiently expressed in *N*. *benthamiana* according to previously described methods (*18*). Briefly, leaves from 4- to 5-week-old plants were infiltrated with suspensions of *Agrobacterium tumefaciens* GV3101 pM90 strains transformed with expression vectors coding for different proteins indicated. Final OD600 of all *A*. *tumefaciens* suspensions were adjusted in infiltration buffer (10 mM MES, 10 mM MgCl2, and 150 μm acetosyringone (pH 5.6)). Final OD600 used was 0.3 for each NLR construct and for each AVRcap1b construct, 0.2 for CP-eGFP, and 0.2 or 0.3 for eGFP depending on whether it was being used as a control for CP or for AVRcap1b, respectively. Whenever multiple constructs were co-infiltrated into an individual spot, the total concentration of bacteria was kept constant across treatments by adding untransformed *A*. *tumefaciens* when necessary. This was to avoid an effect from differences in the total OD600 of bacteria in each treatment.

### Co-immunoprecipitation (CoIP) assays

Co-immunoprecipitation assays were performed as described previously (REF). Four- to five-week-old *N*. *benthamiana* plants were agroinfiltrated as described above with constructs of interest and leaf tissue was collected 3 days post agroinfiltration. Final OD600 used for each construct was 0.3. Leaf tissue was ground using a Geno/Grinder tissue homogenizer. GTEN extraction buffer was used (10% glycerol, 50 mM Tris-HCl (pH 7.5), 5 mM MgCl2, and 50 mM NaCl) supplemented with 10 mM DTT, 1× protease inhibitor cocktail (SIGMA) and 0.2% IGEPAL (SIGMA). Samples were incubated in extraction buffer on ice for 10 min with short vortex mixing every 2 min. Following incubation, samples were centrifuged at 5,000 x*g* for 15 min and the supernatant was collected. This was spun down an additional time at 5,000 *xg* for 15 min and then supernatant was filtered using Minisart 0.45 μm filter (Sartorius Stedim Biotech, Goettingen, Germany).

Part of the extract was set aside prior to immunoprecipitation. These were used as inputs. 1.4 ml of the remaining filtered total protein extract was mixed with 30 μl of anti-FLAG agarose beads (SIGMA) and incubated end over end for 90 min at 4°C. Beads were washed 5 times with immunoprecipitation wash buffer (GTEN extraction buffer with 0.2% v/v IGEPAL (SIGMA)). Associated plant proteins were competitively eluted by excess of 3xFLAG peptides. Elution was spun down at 1,000 *xg* for 1 min and the supernatant was transferred to a new tube. Inputs and eluted immunoprecipitates were diluted in SDS loading dye and denatured by heating for 10 min at 72°C. All samples were used for SDS-PAGE. Briefly, they were run on 4% to 20% Bio-Rad Mini-PROTEAN TGX gels alongside a PageRuler Plus prestained protein ladder (Thermo Scientific). The proteins were then transferred to polyvinylidene difluoride membranes using Trans-Blot Turbo Transfer Buffer using a Trans-Blot Turbo Transfer System (Bio-Rad) as per the manufacturer’s instructions. Membranes were immunoblotted as described below.

### Immunoblotting and detection of SDS-PAGE assays

Blotted membranes were blocked with 5% milk in Tris-buffered saline plus 0.01% Tween 20 (TBS-T) for an hour at room temperature and subsequently incubated with desired antibodies at 4°C overnight. Antibodies used were anti-GFP (B-2) HRP (Santa Cruz Biotechnology), anti-FLAG (M2) HRP (Sigma), or anti-Myc (9E10) HRP (Roche), as indicated in the figures, all used in a 1:5,000 dilution in 5% milk in TBS-T. To visualise proteins, we used Pierce ECL Western (32106, Thermo Fisher Scientific), supplementing with up to 50% SuperSignal West Femto Maximum Sensitivity Substrate (34095, Thermo Fisher Scientific) when necessary. Membrane imaging was carried out with an ImageQuant LAS 4000 or an ImageQuant 800 luminescent imager (GE Healthcare Life Sciences, Piscataway, New Jersey, United States of America). Rubisco loading control was stained using Ponceau S (Sigma) or Ponceau 4R (A.G. Barr).

### Purification of AVRcap1b in complex with NbTOL9a domains

Recombinant AVRcap1b protein (lacking signal peptide and RXLR motif) was expressed by cloning in pOPIN-S3C plasmid, with an N-terminal tandem 6xHis-SUMO followed by a 3C protease cleavage site. pOPIN-S3C:AVRcap1b was transformed into *E. coli* SHuffle cells. Eight litres of these cells were grown at 30°C in autoinduction medium to an OD600 of 0.6 to 0.8 followed by overnight incubation at 18°C and harvested by centrifugation. Pelleted cells were resuspended in 50 mM tris-HCl (pH 8), 500 mM NaCl, 50 mM glycine, 5% (v/v) glycerol, and 20 mM imidazole (buffer A) supplemented with cOmplete EDTA-free protease inhibitor tablets (Roche) and lysed by sonication. The clarified cell lysate was applied to a Ni^2+^-NTA column connected to an AKTA pure system. 6xHis-SUMO-AVRcap1b was step eluted with elution buffer (buffer A with 500 mM imidazole) and directly injected onto a Superdex 200 26/600 gel filtration column pre-equilibrated with buffer B [20 mM Hepes (pH 7.5) and 150 mM NaCl]. The fractions containing 6xHis-SUMO-AVRcap1b were pooled, and the N-terminal 6xHis-SUMO tag was cleaved by addition of 3C protease (10 μg/mg of fusion protein), incubating overnight at 4°C. Cleaved AVRcap1b was further purified using a Ni^2+^-NTA column, this time collecting the flow-through to separate the cleaved tag from the AVRcap1b protein. Untagged AVRcap1b was further purified by another round of gel filtration as described above. The concentration of protein was judged by absorbance at 280 nm (using a calculated molar extinction coefficient of 110,810 M^−1^ cm^−1^ for AVRcap1b).

Recombinant NbTOL9a subdomains, were also expressed cloning in pOPIN-S3C plasmid as described above. NbTOL9a and NbTOL9a^ENTH^ were cloned into pOPIN-S3C plasmids and transformed into *E. coli* sHuffle cells. First, small scale purification trials were performed, expressing 1 litre of each of the 8 transformed *E. coli* strains. Cells were grown at 37°C in autoinduction medium to an OD600 of 0.6 to 0.8 followed by overnight incubation at 18°C and harvested by centrifugation. Pelleted cells were resuspended in 50 mM tris-HCl (pH 8), 500 mM NaCl, 50 mM glycine, 5% (v/v) glycerol, and 20 mM imidazole (buffer A) supplemented with cOmplete EDTA-free protease inhibitor tablets (Roche) and lysed by sonication. The clarified cell lysate was applied to a Ni^2+^-NTA column connected to an AKTA pure system. 6xHis and 6xHis-SUMO-tagged proteins were step eluted with elution buffer (buffer A with 500 mM imidazole), and the elution was used for SDS-PAGE assays.

For scaling up purification of 6xHis-SUMO-NbTOL9a^ENTH^, 8 litres of each *E. coli* SHuffle strain were grown at 37°C in autoinduction medium to an OD600 of 0.6 to 0.8 followed by overnight incubation at 18°C and harvested by centrifugation. Pelleted cells were resuspended in 50 mM tris-HCl (pH 8), 500 mM NaCl, 50 mM glycine, 5% (v/v) glycerol, and 20 mM imidazole (buffer A) supplemented with cOmplete EDTA-free protease inhibitor tablets (Roche) and lysed by sonication. The clarified cell lysate was applied to a Ni^2+^-NTA column connected to an AKTA pure system. 6xHis-SUMO-tagged proteins were step eluted with elution buffer (buffer A with 500 mM imidazole) and directly injected onto a Superdex 75 26/600 gel filtration column pre-equilibrated with buffer B [20 mM Hepes (pH 7.5) and 150 mM NaCl]. The fractions containing 6xHis-SUMO-NbTOL9a^ENTH^ were pooled, and the N-terminal 6xHis-SUMO tag was cleaved by addition of 3C protease (10 μg/mg of fusion protein), incubating overnight at 4°C. Cleaved NbTOL9a^ENTH^ was further purified using a Ni^2+^-NTA column, this time collecting the flow-through to separate the cleaved tag from the protein of interest. Untagged NbTOL9a^ENTH^ was further purified by another round of gel filtration as described above. The concentration of protein was judged by absorbance at 280 nm (using a calculated molar extinction coefficient of 15,470 M^−1^ cm^−1^ for NbTOL9a^ENTH^).

To obtain AVRcap1b in complex with NbTOL9a^ENTH^, both proteins were incubated in a 1:1 molar ratio overnight at 4°C and subjected to gel filtration on a Superdex 200 26/600 gel filtration column as described above. The fractions containing AVRcap1b in complex with NbTOL9a^ENTH-GAT^ or NbTOL9a^ENTH^ were pooled, concentrated to 10 to 15 mg/ml, and subsequently used for crystallization screens.

### Size-exclusion chromatography (SEC) with AVRcap1b and NbTOL9a^ENTH^

AVRcap1b and NbTOL9a were purified as described above. 200 μl of concentrated protein (5 mg/ml) were run on a Superdex 200 Increase 10/300 GL column (Cytiva) connected to an AKTA Pure system (GE Healthcare), with samples being run at a flow rate of 0.4 ml/min. The buffer used for elution was buffer B. Each protein was ran individually first. Then, 200 μl of an equimolar mixture of AVRcap1b and NbTOL9a^ENTH^ was ran using the same procedure, for comparison. The eluted fractions were analysed by SDS–PAGE as described above and stained with Coomassie.

### Size-exclusion chromatography (SEC)-coupled multi-angle light scattering (MALS)

SEC-MALS was performed using a Superdex 200 Increase 10/300 GL column (Cytiva) connected to a DAWN multi-angle light scattering detector (Wyatt Technology, Santa Barbara, CA), with an inline Optilab refractive index detector. 500 µg of purified protein was injected and separated at a flow rate of 0.5 mL/min in 20 mM HEPES (pH 7.5), 150 mM NaCl. For the AVRcap1-NbTOL9a^ENTH^ complex, proteins were mixed at equimolar concentrations and incubated on ice prior to injection. Molecular weight calculations were performed across the elution peak using ASTRA software (Wyatt Technology), assuming a refractive index increment (dn/dc) of 0.186 mL/g, appropriate for unmodified proteins (*59*).

### Crystallization, data collection and structure solution of AVRcap1b-NbTOL9a^ENTH^ protein complex

Crystallisation screens were performed at 21°C using the sitting-drop vapour diffusion technique. Drops composed of 0.3 μL of protein solution and 0.3 μL of reservoir solution were set up in MRC 96-well crystallisation plates (Molecular Dimensions), which were dispensed using an Oryx Nano or an Oryx8 robot (Douglas Instruments). Crystal growth was monitored using a Minstrel Desktop Crystal Imaging System (Rikagu). Suitable crystals grew from a BCS screen crystallisation condition containing 15% (v/v) PEG Smear Medium, 5% (v/v) 2-propanol, 10% (v/v) ethylene glycol, 0.2 M magnesium chloride hexahydrate in 0.1 M HEPES buffer pH 7.5 (Molecular Dimensions) and were harvested after the addition of 30% (v/v) glycerol by flash-cooling in liquid nitrogen using LithoLoops (Molecular Dimensions). X-ray diffraction data were collected at the Diamond Light Source (Didcot, UK) on beamline I04 using an Eiger2 XE 16M pixel array detector (Dectris) with crystals maintained at 100 K by a Cryojet cryocooler (Oxford Instruments).

X-ray data were integrated, and scaled using DIALS (*60*) as implemented through the XIA2 (*60*) pipeline, and then merged using AIMLESS (*60*), via the CCP4i2 graphical user interface (*61*). The AVRcap1b-ENTH complex crystallised in space group *P*212121 with cell parameters *a*=85.9, *b*=136.9, *c*=195.6 Å, and the best crystal yielded diffraction data to only 4.1 Å resolution (see **Table S1** for a summary of data collection and processing statistics). Estimation of the asymmetric unit (ASU) composition suggested two copies of a 1:1 complex giving a 63% solvent content.

AlphaFold2-multimer (*62*), as implemented through Colabfold (https://colab.research.google.com/github/sokrypton/ColabFold/blob/main/AlphaFold2.ipyn <u>b)</u> (*63*) was used to generate templates for molecular replacement. There was very good sequence coverage for both proteins and the five independent models of the individual components were closely similar. The pLDDT scores were generally good (averages of ≥86 for AVRcap1b and ≥78 for ENTH models). However, the relative placement of the two components of the complex varied across the five models and the corresponding PAE scores indicated very low confidence in these predictions and, therefore, none of these were appropriate inputs for molecular replacement without further processing. Templates were prepared from the rank 1 AlphaFold2 model using the phenix.process_predicted_model tool within the Phenix package (*64*) to correct *B*-factors, remove low confidence regions and divide the model into segments as appropriate. Using an RMSD cutoff of 1.5 Å, AVRcap1b was split into three segments and ENTH remained as a single domain, thereby giving a total of four separate input templates. PHASER (*65*) was able to place two copies of each template, apart from one template from AVRcap1b, where only a single copy was located. After rearranging these into coherent assemblies in COOT (*66*), the missing segment could be placed into vacant density guided by the arrangement of the segments within the more complete copy of AVRcap1b. The structure was then subjected to jelly body refinement in REFMAC5 (*67*) using ProSMART restraints (*68*) generated from the starting AlphaFold2 model, giving *R*work and *R*free values of 0.360 and 0.376, respectively, to 4.1 Å resolution. Now it was possible to generate more complete models for the components by superposing the original unprocessed AlphaFold2 model and trimming this with reference to the improved electron density. Due to the low resolution of the dataset, only very limited rebuilding was possible in COOT, where Geman-McClure and Ramachandran restraints were used to maintain good stereochemical parameters. After several cycles of restrained refinement in REFMAC5 with TLS restraints (with each protein chain treated as a separate TLS domain), and editing in COOT, the final model was obtained with *R*work and *R*free values of 0.248 and 0.286, respectively to 4.1 Å resolution (see **Table S1** for a summary of refinement statistics). All structural figures were prepared using ChimeraX (*69*).

### Phylogenetic analysis

We used the *Phytophthora infestans* AVRcap1b protein sequence as a query in a PSI-BLAST search (https://blast.ncbi.nlm.nih.gov/Blast.cgi) against the NCBI non-redundant protein database (*1*). After the first iteration, we observed that PSR2, a phylogenetically distant effector from Phytophthora sojae, appeared among the top 250 hits (**Data S1**). Based on this observation, we terminated the search after one round and retained the top 250 hits. To focus on sequences similar in size to AVRcap1b, we filtered out those shorter than 500 amino acids or longer than 800 amino acids, resulting in a final set of 180 sequences (**Data S2, Table S4**). We then supplemented this dataset with AVRcap1b and three of its orthologs from *P. andina*, *P. mirabilis*, and *P. ipomoeae*. Sequences were aligned using MAFFT v7.526 with the options [--anysymbol --localpair] (*9*). he resulting alignment was used to construct a phylogenetic tree with FastTree v2.1.11 using the LG substitution model (option -lg) (*10*). Phylogenetic trees were visualized with iTOL v6 (*11*). A tree of AVRcap1b clade was generated using the same method. Phylogenetic trees and sequence alignments can be found on the Zenodo repository (*70*). All scripts are deposited at [https://github.com/amiralito/AVRcap1b] repository.

### Structural predictions

The regions downstream of the RXLR motif (after the ‘DEER’ motif) from representative sequences in the AVRcap1b-like and PSR2-like clades were extracted based on alignment with AVRcap1b and modeled using AlphaFold 3 with seed = 1 (*46*). The top-ranked model for each prediction was colored according to pLDDT values using data from the AlphaFold summary JSON file and visualized in ChimeraX (*69*). Structural alignments were performed with the matchmaker command in ChimeraX, which was also used to generate the structure figures. All predicted models are deposited on Zenodo repository (*70*).

### Shannon’s Entropy and conservation analysis

Sequences from the AVRcap1b-like clade were filtered by length to retain only those between 550 and 750 amino acids, resulting in 128 sequences. Redundant sequences were removed using CD-HIT with parameters [-c 1.0 -d 100], yielding 120 unique sequences (*71*). These sequences were aligned using MAFFT v7.526 with the options [--anysymbol --localpair] (*72*), and the resulting alignment was trimmed with ClipKIT v2.3.0 using the [-l -m gappy] settings to remove poorly aligned and gapped regions (*73*). The final trimmed alignment was used to calculate Shannon’s entropy per position using the ‘entropy’ R package (*74*), and to generate sequence logos with the ‘ggseqlogo’ and ‘ggplot2’ R packages (*75, 76*), as part of the same approach reported before (*16, 21*), following an approach previously described (**Table S3**) (*50*). AVRcap1b phylogenetic subclades were defined based on well-supported branches defining the clade and Phytophthora clades present in each subclade and were extracted using Dendroscope v3.8.1 [Options > Advanced Options > Extract Subnetwork] (*77*). All scripts are available at [https://github.com/amiralito/AVRcap1b].

## Acknowledgements

We thank J. Upson (The Sainsbury Laboratory, Norwich, UK) for assistance with generating constructs for expression of clade 1c AVRcap1b orthologs. We thank J. Mundy (John Innes Cenre) for assistance with crystallography experiments. We thank the Diamond Light Source, UK (beamline i04 under proposal MX25108) for access to x-ray data collection facilities. J.M., A.T., M.P.C and S.K. thank Alf A. Fold for invaluable assistance and support. M.P.C. thanks D. C. Gómez De La Cruz (The Sainsbury Laboratory) for support throughout the project. We thank all members of the TSL Support Services for their invaluable assistance.

## Author contributions

Conceptualization: J.M., A.T., J.C.D.L.C., L.D., T.O.B., M.J.B., S.K. and M.P.C.

Methodology: J.M., A.T., A.R.B., J.C.D.L.C., D.M.L., C.E.M.S., A.V.C., T.O.B., and M.P.C.

Data curation: J.M., A.T., D.M.L., and M.P.C.

Formal analysis: J.M., A.T., D.M.L., and M.P.C.

Investigation: J.M., A.T., H.P, M.H., A.R.B., E.L.H.Y., D.M.L., C.E.M.S., A.V.C., L.D., and M.P.C.

Resources: D.M.L., C.E.M.S., L.D., T.O.B., M.J.B., S.K. and M.P.C.

Writing—original draft: J.M., A.T., A.R.B., B.A.S., D.M.L., S.K. and M.P.C.

Writing—review and editing: TBD.

Visualization: J.M., A.T., A.R.B., B.A.S., M.P.C.

Supervision: J.M., M.J.B., S.K. and M.P.C.

Project administration: J.M., A.T., S.K. and M.P.C.

Funding acquisition: S.K.

## Data and materials availability

All data needed to evaluate the conclusions in the paper are present in the paper and/or the Supplementary Materials. Data related to the AVRcap1b-NbTOL9a^ENTH^ structure can be found at PDB, with PDB ID 9RDC, as well as in **Table S1**. Datasets associated to AlphaFold 3 effector predictions can be accessed on Zenodo (*70*). All scripts used to extract information from AlphaFold predicted structures and to generate the entropy plots are available at https://github.com/amiralito/AVRcap1b.

## Supplementary Materials

**Figure S1.**
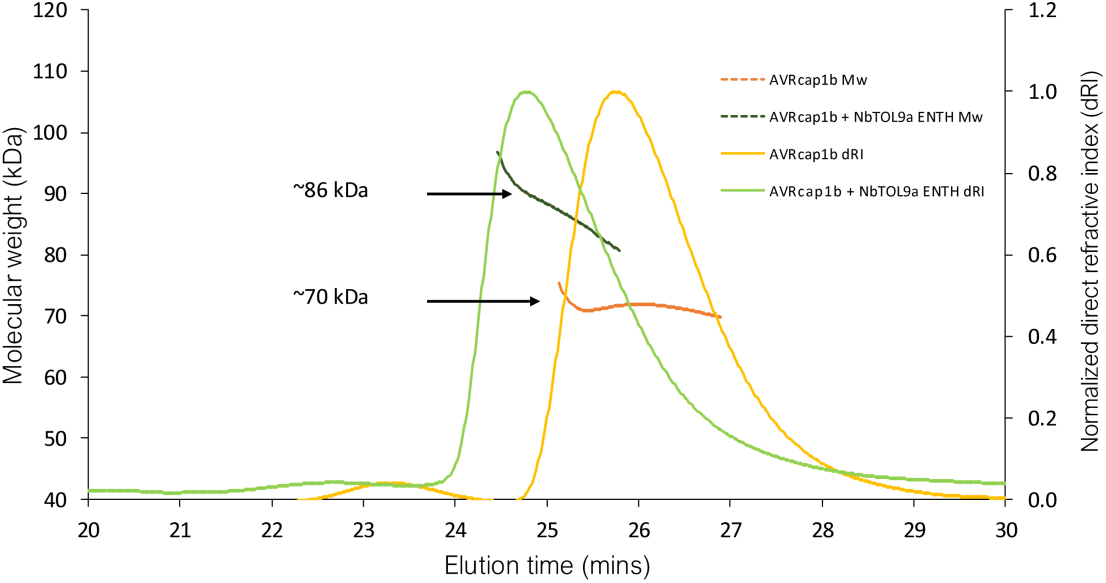
SEC-MALS confirms complex formation between AVRcap1b and NbTOL9a^ENTH^. Size-exclusion chromatography coupled to multi-angle light scattering (SEC-MALS) was used to determine the molar mass of AVRcap1b alone (yellow) and in complex with NbTOL9a^ENTH^ (green). Normalized differential refractive index (dRI) traces represent elution profiles from a Superdex 200 Increase 10/300 GL column. Dashed lines indicate the calculated molecular weight (kDa) across the elution peaks. Incubation of AVRcap1b with NbTOL9a^ENTH^ resulted in a shift in elution volume and an increase in calculated molecular weight from ∼70 kDa (AVRcap1b alone) to ∼86 kDa, consistent with the formation of a 1:1 complex in solution.

**Figure S2:**
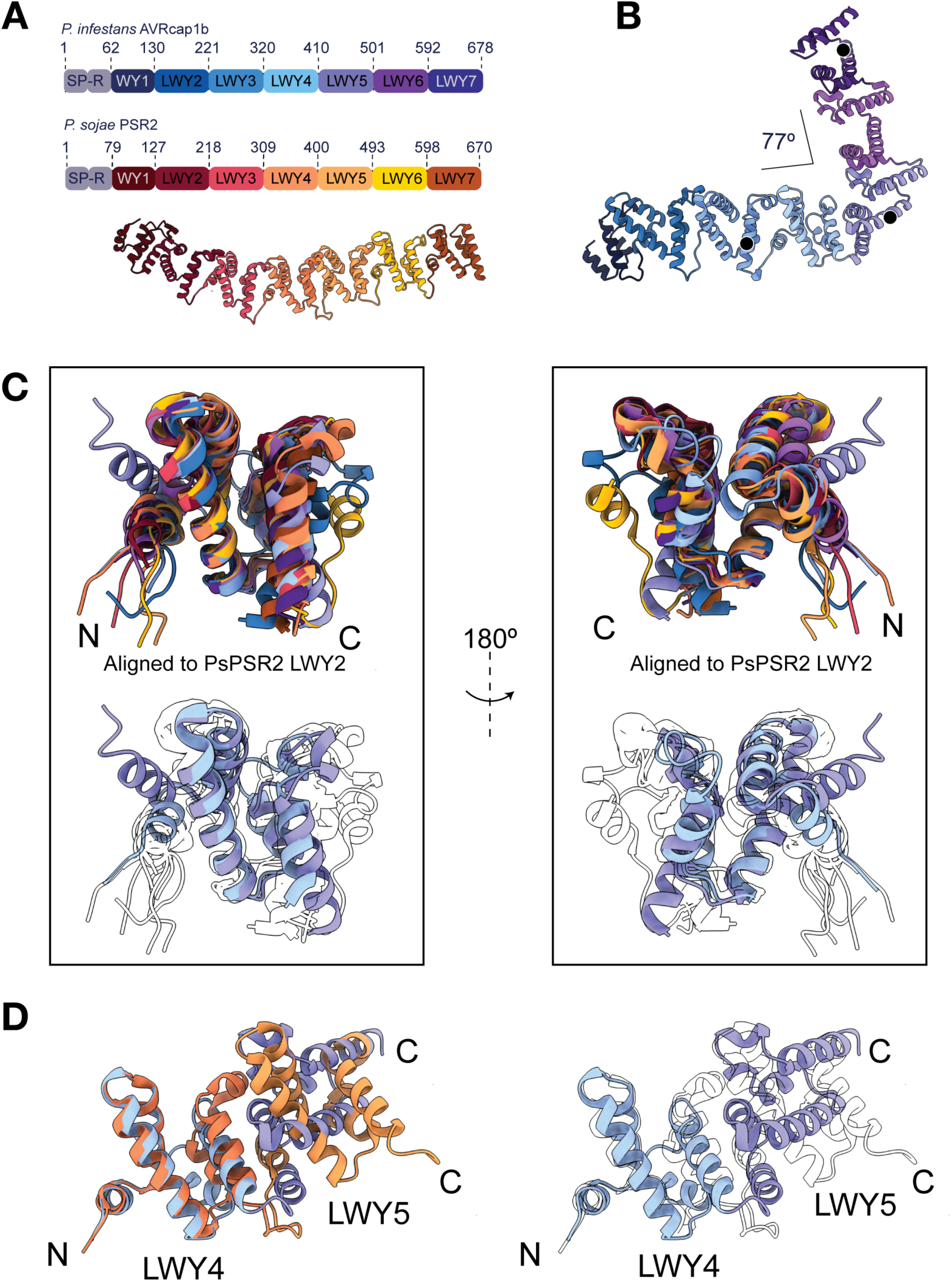
LWY4 and LWY5 of AVRcap1b are structurally distinct from other LWY modules. (**A**) Schematic representation of *P. infestans* AVRcap1b and *P. sojae* PSR2. Individual structural repeat units of each effector are color-coded as indicated in the schematic, which also denotes the amino acid boundaries for each repeat/domain. The schematic highlights the signal peptide (SP) and RXLR-DEER motif (R) at the N-terminus. Overall structure of *P. sojae* PSR2 is also included (PDB ID: 5GNC). (**B**) Overall L-shaped structure of AVRcap1b, highlighting the 77° angle introduced by LWY4 and LWY5 repeats. Black dots indicate amino acids selected to calculate the angle. (**C**) Two different views of a structural alignment of all LWY repeats from PSR2 and AVRcap1b. Bottom images feature the same alignment with all repeats made transparent and LWY4 and LWY5 from AVRcap1b in color. (**D**) Structural alignment of LWY4 and LWY5 of PSR2 and AVRcap1b. Color coding is as indicated in the schematics in panel (**A**).

**Figure S3:**
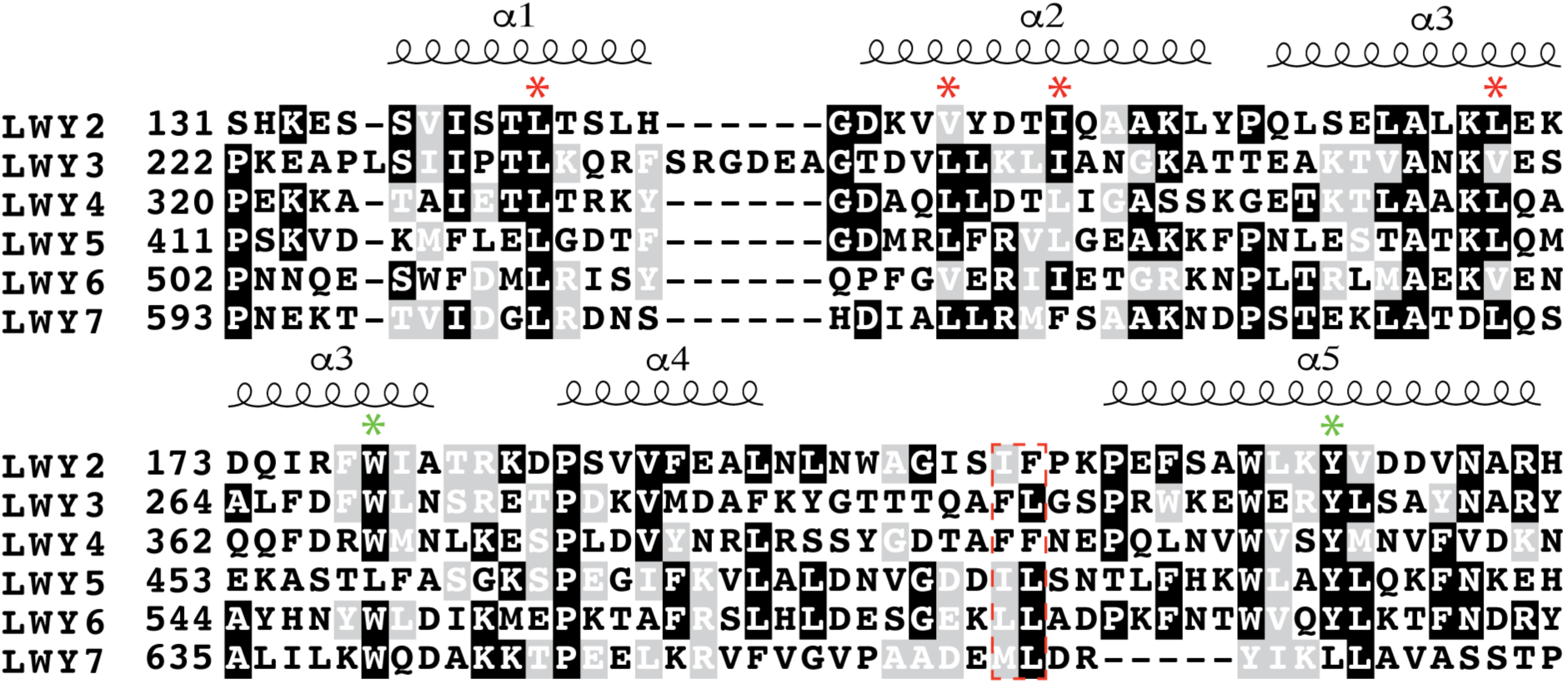
Amino acid sequence alignment of all LWY modules of AVRcap1b. The conserved hydrophobic Loop^4-5^ is highlighted with a red dashed line. The hydrophobic residues that make up the conserved hydrophobic pocket in each LWY module are shown with red asterisks. Green asterisks highlight the conserved W and Y residues. Secondary structure is shown at the top.

**Figure S4.**
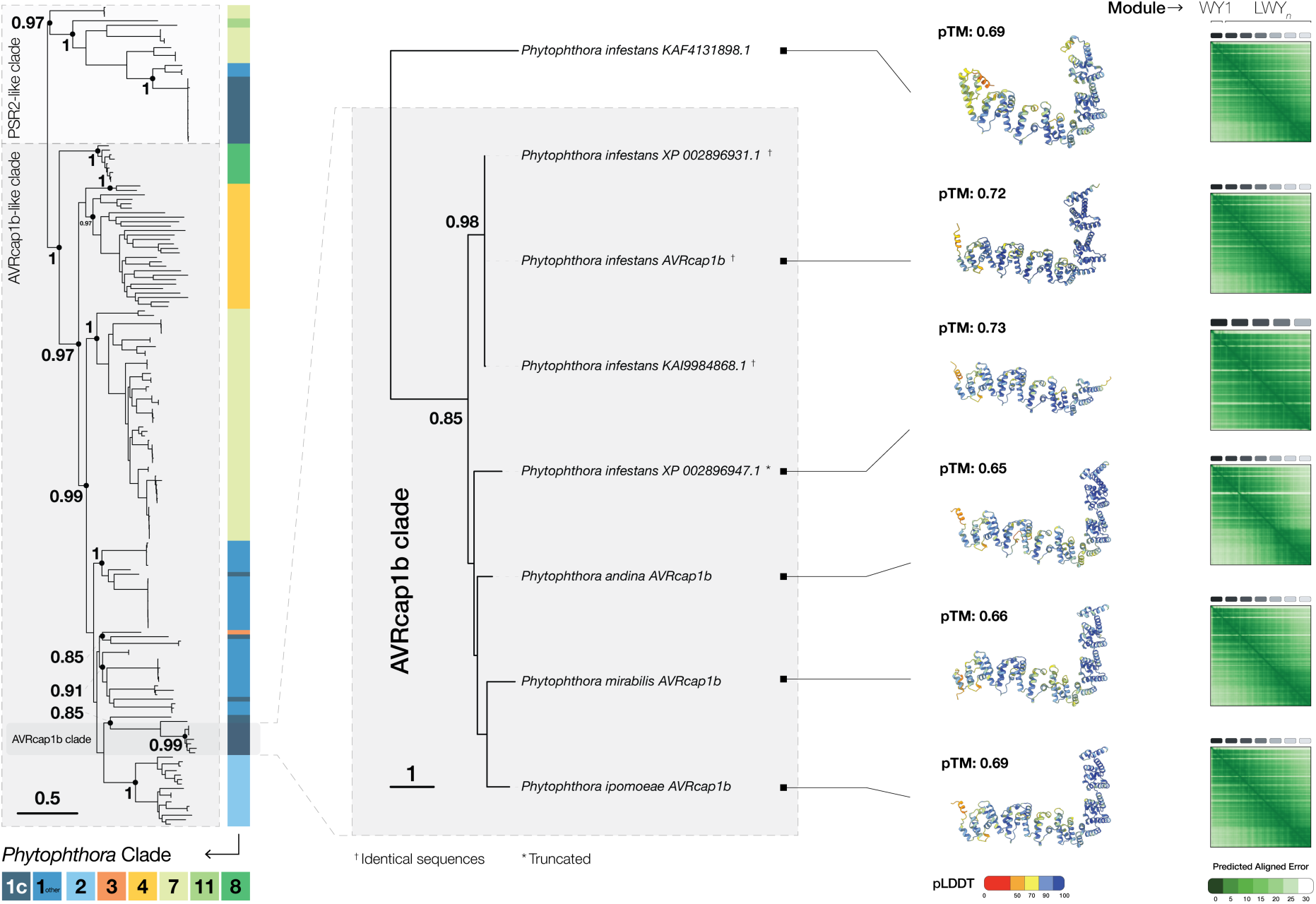
Clade 1c AVRcap1b orthologs all share the same L-shaped effector fold. A magnified view of the AVRcap1b subclade from the phylogenetic tree of AVRcap1b-like and PSR2-like sequences shows that all clade 1c orthologs adopt the characteristic L-shaped fold, consistent with other AVRcap1b-like effectors. One exception is the *P. infestans* sequence XP_002896947.1, which was predicted to have a shorter, stick-like structure due to the absence of LWY6 and LWY7 repeats. Structural modeling was performed using AlphaFold 3 (*46*).

**Figure S5.**
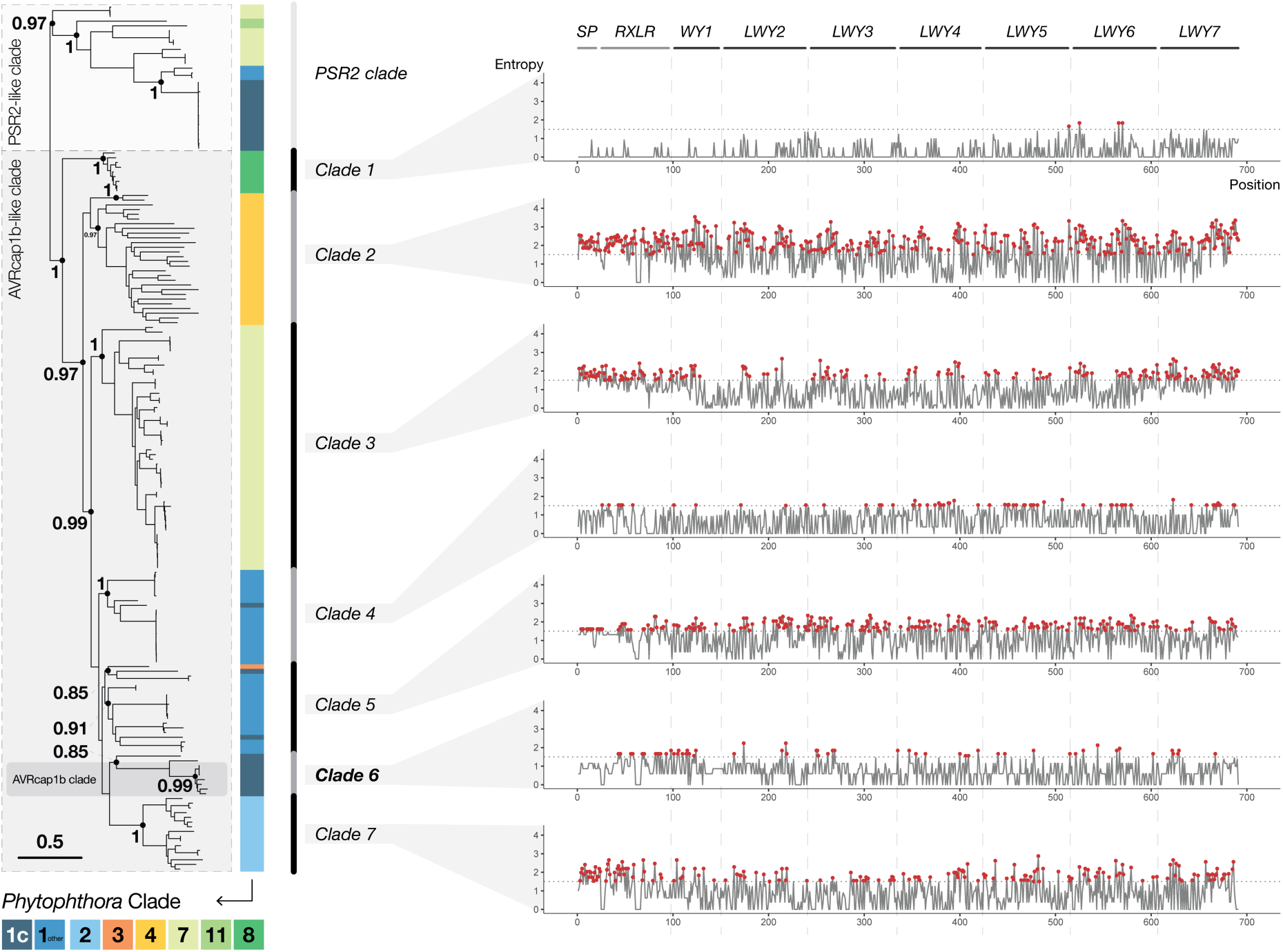
AVRcap1b subclades exhibit variable Shannon’s entropy patterns. Entropy was calculated across AVRcap1b-like sequences grouped into subclades. Sites with entropy values >1.5 were considered highly variable. WY and LWY modules exhibited extensive variation both within and between clades. In clade 6, which includes AVRcap1b and its clade 1c orthologs, WY1 was the most variable module, with 10 highly variable sites compared to other LWY modules (**Table S2**).

**Figure S6:**
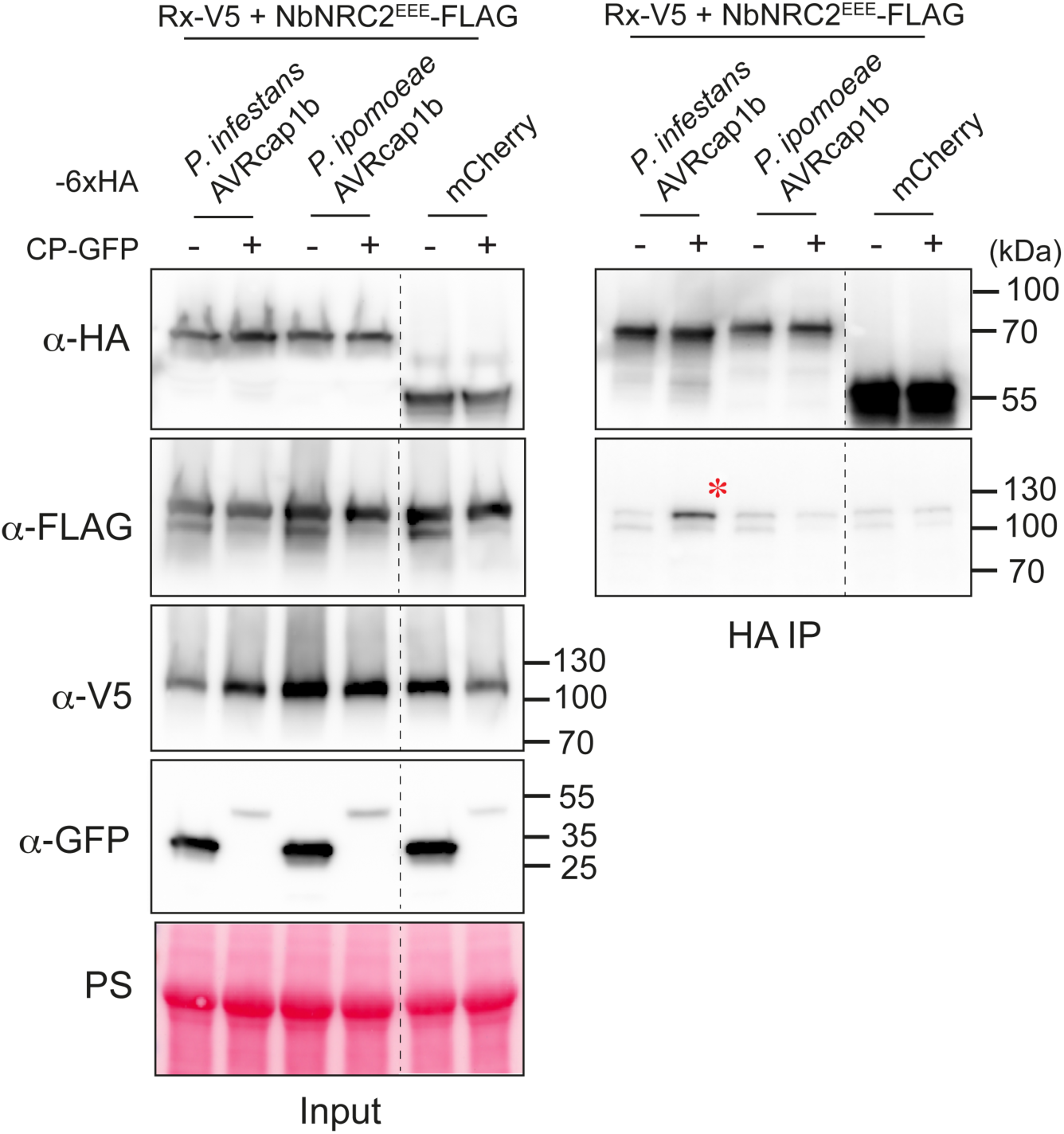
Unlike *P. infestans* AVRcap1b, the *P. ipomoeae* ortholog does not associate with Rx/CP-activated NbNRC2. CoIP assays between resting state or activated NbNRC2 and AVRcap1b variants. C-terminally 3xFLAG-tagged NbNRC2 was coexpressed with C-terminally V5-tagged Rx and C-terminally 6xHA-tagged AVRcap1b. the Rx/NbNRC2 system was activated with C-terminally GFP tagged PVX CP. Free GFP used as a negative control for activation. IPs were performed with agarose beads conjugated to HA antibodies (HA IP). Total protein extracts were immunoblotted with the antisera labeled on the left. Approximate molecular weights (in kilodalton) of the proteins are shown on the right. Red asterisk indicates AVRcap1b association with Rx/CP activated NbNRC2^EEE^. Rubisco loading control was carried out using Ponceau stain (PS). The experiment was repeated three times with similar results.

**Table S1:** Summary of X-ray data and model parameters for AVRcap1b – NbTOL9a^ENTH^ complex.

|  |  |
| --- | --- |
| **Data collection** |  |
| Diamond Light Source beamline | I04 |
| Wavelength (Å) | 0.9795 |
| Detector | Eiger2 XE 16M |
| Resolution range (Å) | 79.57 – 4.10 (4.49 – 4.10) |
| Space Group | *P*2<sub>1</sub>2<sub>1</sub>2<sub>1</sub> |
| Cell parameters (Å) | *a* = 85.9, *b* = 136.9, *c* = 195.6 |
| Total no. of measured intensities | 242580 (59716) |
| Unique reflections | 18772 (4401) |
| Multiplicity | 12.9 (13.6) |
| Mean *I*/$\sigma$(*I*) | 9.6 (1.0) |
| Completeness (%) | 100.0 (100.0) |
| *R*<sub>merge</sub><sup>a</sup> | 0.106 (2.126) |
| *R*<sub>meas</sub><sup>b</sup> | 0.110 (2.208) |
| *CC*<sub>1/2</sub><sup>c</sup> | 1.000 (0.518) |
| **Refinement** |  |
| Resolution range (Å) | 79.57 – 4.10 (4.21 – 4.10) |
| Reflections: working/free<sup>d</sup> | 17765/938 |
| *R*<sub>work</sub>/*R*<sub>free</sub><sup>e</sup> | 0.248/0.286 |
| MolProbity score/Clashscore<sup>f</sup> | 1.39/0.97 |
| Ramachandran plot: favoured/allowed/disallowed<sup>f</sup> (%) | 98.2/1.8/0.0 |
| R.m.s. bond distance deviation (Å) | 0.005 |
| R.m.s. bond angle deviation (°) | 1.53 |
| AVRcap1b – chains/residue ranges | A,B/78-675,78-675 |
| ENTH – chains/residue ranges | C,D/1-138,3-138 |
| PDB accession code | 9RDC |

Values in parentheses are for the outer resolution shell.
<sup>a</sup> $R_{\text{merge}} = \sum_{hkl} \sum_i |I_i(hkl) - \langle I(hkl) \rangle| / \sum_{hkl} \sum_i I_i(hkl)$ .
<sup>b</sup> $R_{\text{meas}} = \sum_{hkl} [N/(N-1)]^{1/2} \times \sum_i |I_i(hkl) - \langle I(hkl) \rangle| / \sum_{hkl} \sum_i I_i(hkl)$ , where $I_i(hkl)$ is the *i*th observation of reflection *hkl*, $\langle I(hkl) \rangle$ is the weighted average intensity for all observations *i* of reflection *hkl* and *N* is the number of observations of reflection *hkl*.
<sup>c</sup> *CC*<sub>1/2</sub> is the correlation coefficient between symmetry equivalent intensities from random halves of the dataset.
<sup>d</sup> The data set was split into "working" and "free" sets consisting of 95 and 5% of the data respectively. The free set was not used for refinement.
<sup>e</sup> The R-factors *R*<sub>work</sub> and *R*<sub>free</sub> are calculated as follows: $R = \sum(|F_{\text{obs}} - F_{\text{calc}}|) / \sum |F_{\text{obs}}|$ , where *F*<sub>obs</sub> and *F*<sub>calc</sub> are the observed and calculated structure factor amplitudes, respectively.
<sup>f</sup> As calculated using MolProbity (78).

**Table S2: RMSDs of pairwise structural comparisons between LWY domains of AVRcap1b and PSR2. Provided as a separate file.**

(**A**) Pairwise comparisons of all LWY domains of *P. infestans* AVRcap1b. (**B**) Pairwise comparisons of all LWY domains of *P. infestans* AVRcap1b to all LWY domains of *P. sojae* PSR2. Columns and rows highlighted in orange indicate comparisons that involve LWY5 from *P. infestans* AVRcap1b.

**Table S3:** Shannon’s Entropy values for AVRcap1b subclades. Provided as a separate file.

**Table S4:** Metadata of AVRcap1b and PSR2 orthologs used for the phylogenetic analysis. Provided as a separate file.

**Data S1:** PSI-BLAST output of AVRcap1b search against NCBI non-redundant protein database. Provided as a separate file.

**Data S2:** Fasta sequences of AVRcap1b and PSR2 orthologs used for the phylogenetic analysis. Provided as a separate file.

**Data S3:** Fasta sequences of Phytophthora clade 1c AVRcap1b sequences used in this study. Sequences provided are for the mature protein, without including the N-terminal signal peptide or RXLR-DEER motif. Provided as a separate file.

**Data S4:** Fasta sequences of AVRcap1b sequences used for Shannon’s Entropy analysis. Provided as a separate file.

